# Large-scale high-density brain-wide neural recording in nonhuman primates

**DOI:** 10.1101/2023.02.01.526664

**Authors:** Eric M. Trautmann, Janis K. Hesse, Gabriel M. Stine, Ruobing Xia, Shude Zhu, Daniel J. O’Shea, Bill Karsh, Jennifer Colonell, Frank F. Lanfranchi, Saurabh Vyas, Andrew Zimnik, Natalie A. Steinmann, Daniel A. Wagenaar, Alexandru Andrei, Carolina Mora Lopez, John O’Callaghan, Jan Putzeys, Bogdan C. Raducanu, Marleen Welkenhuysen, Mark Churchland, Tirin Moore, Michael Shadlen, Krishna Shenoy, Doris Tsao, Barundeb Dutta, Timothy Harris

## Abstract

High-density, integrated silicon electrodes have begun to transform systems neuroscience, by enabling large-scale neural population recordings with single cell resolution. Existing technologies, however, have provided limited functionality in nonhuman primate species such as macaques, which offer close models of human cognition and behavior. Here, we report the design, fabrication, and performance of Neuropixels 1.0-NHP, a high channel count linear electrode array designed to enable large-scale simultaneous recording in superficial and deep structures within the macaque or other large animal brain. These devices were fabricated in two versions: 4416 electrodes along a 45 mm shank, and 2496 along a 25 mm shank. For both versions, users can programmatically select 384 channels, enabling simultaneous multi-area recording with a single probe. We demonstrate recording from over 3000 single neurons within a session, and simultaneous recordings from over 1000 neurons using multiple probes. This technology represents a significant increase in recording access and scalability relative to existing technologies, and enables new classes of experiments involving fine-grained electrophysiological characterization of brain areas, functional connectivity between cells, and simultaneous brain-wide recording at scale.

## Introduction

High-channel count electrophysiological recording devices such as Neuropixels probes (*1, 2*) are transforming neuroscience with rodent models by enabling recording from large populations of neurons anywhere in the rodent brain (*3–9*). The capabilities provided by this approach have led to important new discoveries, such as a continuous attractor network of grid cells in the rat hippocampus (*9*), establishing a causal topographical mapping between activity in the cortex and striatum in mice (*4*), and revealing how thirst modulates brain-wide neural population dynamics in mice (*7*). The Neuropixels 1.0 probe has also been used to record neurons acutely in nonhuman primates (NHPs) like macaques (*10–13*) and in humans (*14, 15*), but their short length (10 mm) restricts access to all but the most superficial targets, and the thinness of the shanks (24 µm) renders them fragile and difficult to insert through primate dura mater.

In a number of discussions with a community of primate researchers, scientists articulated the need for a probe more suitable for use with rhesus macaques. In particular, this community desired a technology that is capable of easily accessing as many neurons in as much of the brain as possible, and a technology which is compatible with conventional acute (single-session) recording techniques. These needs collectively suggested a probe design with a long linear array of dense recording sites, similar to the rodent probes, but with mechanical properties that would enable transdural insertion in macaques.

Current commercially-available technologies do not fully satisfy these needs. Linear array technologies, such as V or S-probes (Plexon Inc.), provide access to the whole brain, but are limited to 64 channels and have relatively large diameters (e.g., 380 µm) that increase as recording channels are added. Surface arrays like the Utah array (*16*) or floating microwire arrays (*17*), allow for recording from up to 256 channels simultaneously, but are limited to recording at pre-specified depths in the superficial cortex, require opening the dura for placement, and cannot be moved after implantation. This latter constraint adds considerable risk to an experimental workflow, as such surgeries may not yield successful recording quality in animals trained for years on an experimental task. Alternative approaches using individually-driven single electrodes have achieved recordings in deep brain regions from dozens or hundreds of neurons by a few dedicated labs (*18–20*), though these approaches do not allow for dense sampling within a specific target region.

Alternatively, two photon (2P) calcium imaging, allows for large-scale population recordings at single cell resolution, but with limited temporal resolution (due to both calcium signaling kinetics and imaging scan rates) and limited recording depth (0–500 µm from the pial surface). In addition, the genetic modification required for 2P imaging remains challenging to implement reliably (*21*). While approaches like microendoscopic imaging have been developed to address the imaging depth limitation, these require insertion of large (1mm) imaging lenses and provide access to a comparably small number of cells without moving the implanted lens (*22*).

We developed the Neuropixels 1.0-NHP, a high-density integrated silicon electrode array, optimized for recording in NHPs, and designed to enable flexible and configurable recording from large populations of neurons throughout the entire brain with single-neuron and single-spike resolution. While the probe’s design and electronic specifications are based on the Neuropixels 1.0 probe (http://Neuropixels.org), fabricating these probes with the desired combination of mechanical and electrical properties using a standard CMOS photolithography process required significant engineering advances. Each probe is larger than a photolithographic reticle, requiring “stitching” of electrical traces across the boundaries between multiple reticles in order to achieve the required probe geometry (*23*) and the introduction of stress compensation layers to prevent intrinsic material stresses from causing the shank to bend. Relative to Neuropixels 1.0, the two variants of Neuropixels-NHP feature a longer, wider, and thicker shank (45 mm long, 125 µm wide, and 90 µm thick and 25 mm long, 125 µm wide, and 60 µm thick; Fig. 1A). For each probe variant, the full length of the shank is populated with recording sites with a density of 2 sites every 20 micrometers (Fig. 1B). A switch under every site allows flexible selection of the 384 simultaneous recording channels across these 4416 or 2496 sites (11 or 6 banks of 384 channels respectively plus one half-sized bank at the shank-base junction; Fig. 1C, Supplementary Fig. S1).

**Figure 1.**
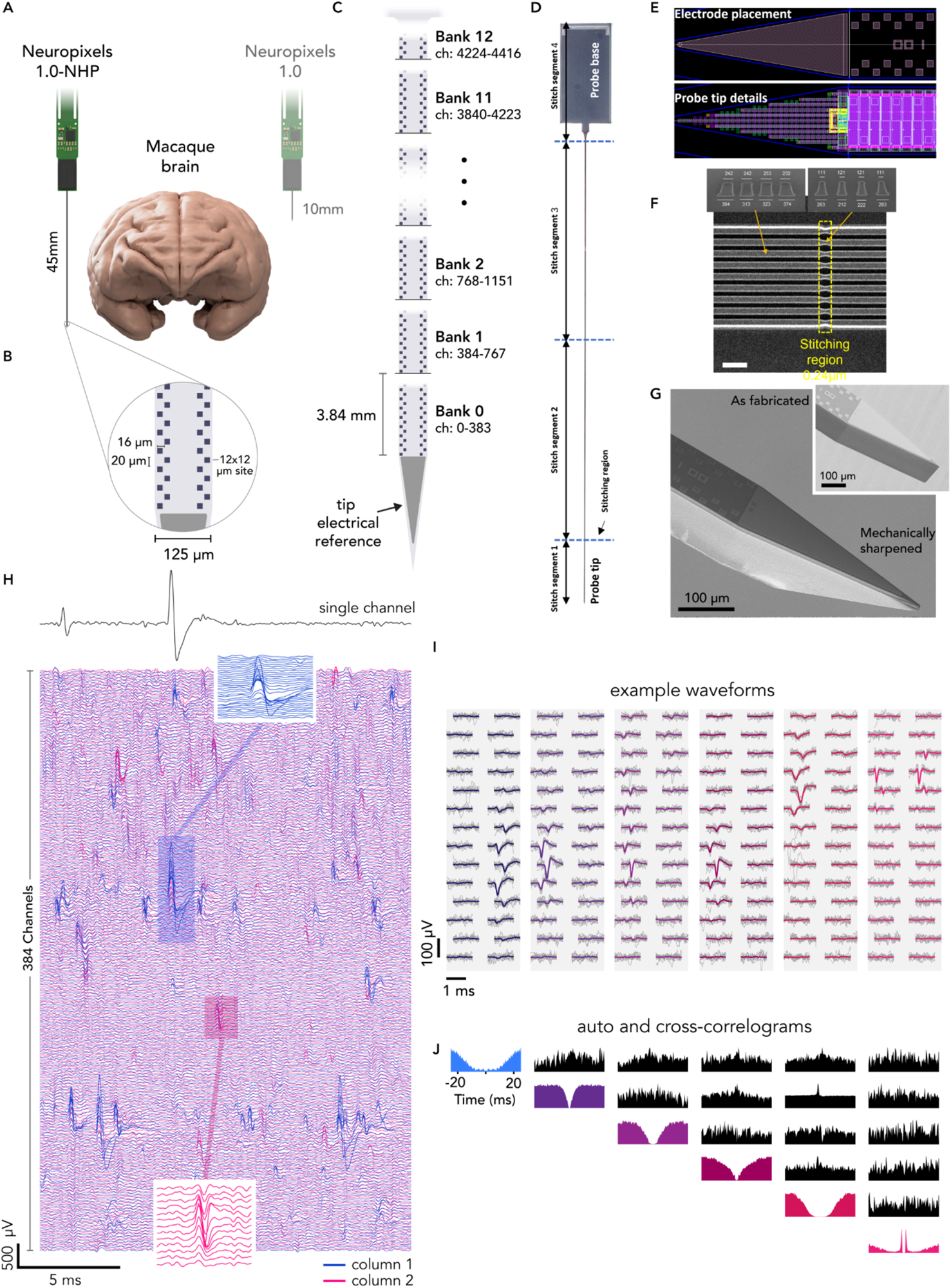
Neuropixels 1.0-NHP probe characterization and engineering. **(A)** comparison of probe geometry with macaque brain and Neuropixels 1.0 probe. **(B)** electrode site layout. **(C)** 4416 recording sites cover the full length of the shank, grouped in 11.5 banks of 384 channels. **(D)** Neuropixels 1.0-NHP probe silicon die photograph indicating the four segments or subblocks used for the sub-field stitching fabrication process. **(E)** Details of the shank tip electrode layout (top) and CMOS circuit layout (bottom). **(F)** Top-down scanning-electron-microscope (SEM) image of one of the metal layers in the shank when crossing the stitching region (scale bar: 1 µm); top-left: cross-section taken outside the stitching overlap region; top-right: cross-section at the narrowest point; the metal wires are narrower due to the double resist exposure. **(G)** SEM picture of shank tip mechanically ground out-of-plane to 25°; inset: probe tip geometry as fabricated, pre-grinding. **(H)** raw electrical recordings from 384 simultaneous channels in the motor cortex of a rhesus macaque monkey. Insets: expanded view of single spike events from single neurons. **(I)** Example waveforms from isolated single neurons (gray) and median waveform (colored). **(J)** auto and cross correlograms for neurons shown in (I).

Here, we describe this novel probe, the engineering challenges surmounted in fabrication, and illustrate the ability to address a range of novel experimental use cases for neuroscientific data collection in large animal models. In particular, we focus on recording large populations of neurons, recording deep in the brain, and recording neurons from multiple brain regions simultaneously using one or multiple probes. We illustrate these use cases with example experiments designed to address specific questions in sensory, motor, and cognitive neuroscience using macaques. The scale and access provided by Neuropixels 1.0-NHP enable a wide range of new experimental paradigms, while streamlining neural data collection at a fraction of the cost of existing alternatives.

## Results

### Technology

The Neuropixels 1.0-NHP probe uses the same signal-conditioning circuits as the Neuropixels 1.0 probe (*1*), integrating 384 low-noise readout channels with programmable gain and 10-bit resolution, in a 130nm Silicon-on-Insulator CMOS platform. The probe shank consists of an array of shift-register elements and a switch matrix to enable the selection of electrode groups. The probe is fabricated as a monolithic piece of silicon, which includes the shank and base electronics, and the total probe lengths are 54 mm and 34 mm for the longer and shorter versions respectively, both larger than the optical reticle. Methods, known as *stitching*, were developed to overcome this limitation (*24*), which involves aligning features from different exposures between adjacent reticles.

The typical maximum size of a 130 nm CMOS chip is limited by the maximum reticle size (ie., 22 mm x 24 mm in this case) that can be exposed in a single lithography step. To build larger chips, the CMOS chip is divided into sub-blocks smaller than the reticle field, which are later used to re-compose (i.e. stitch together) the complete chip by performing multiple reticle exposures per layer. The stitched boundary regions of the small design blocks are overlapped (i.e., double exposed) to ensure uninterrupted metal traces from one reticle to the next, which is accounted for in the design. Since multiple levels of metal wires are needed to realize and interconnect the large number of electrodes at high density, the conducting wires in the shank must be aligned between adjacent reticles for four of the six stacked aluminum metal layers. To enable stitching and to simplify photo-mask design and reuse, the probe is designed with three elements as shown in Fig. 1D: a 5 mm tip (Fig. 1E), either one or two 20 mm middle shank segments (stitch segments 2 and 3 in Fig. 1D) for the 25 mm and 45 mm versions respectively, and the 10 mm base segment. Thus either three reticles (25 mm shanks) or four reticles (45 mm shanks) are required for each probe. The probe base is 6 mm wide, allowing four probes to be written across these three (25 mm shank) or four (45 mm shank) reticles.

Four design strategies were developed to mitigate the expected degradation of signal with increased shank length: 1) the 384 metal wires connecting the electrodes to the recording circuits in the base were made wider than the Neuropixels 1.0 probe shank to keep their resistance and thermal noise contribution low; 2) the spacing between the metal wires was increased to limit the signal coupling and crosstalk along the shank; 3) the power-supply wires connected to shank circuits were made wider to minimize voltage attenuation and fluctuations along the shank; and 4) the size of the decoupling capacitors is increased. The additional width and spacing of the metal lines also mitigated the impact of anticipated reticle misalignments, magnification, and rotational errors in the overlap regions during stitching. Fig. 1E shows details of the shank electrode placement and the dense CMOS layout in the shank and Fig. 1F shows a scanning-electron-microscope (SEM) image of the Al metal wires running along one of the 0.24-µm stitching regions in the shank. Due to the double exposure of the masking photoresist at the stitching region, the wires become 24%-54% narrower in this region, but remain continuous circuits without interruption.

One of the major design changes to this device, compared to the Neuropixels 1.0 (rodent) probe (*1*), was to strengthen and thicken the probe shank, which was necessary both to support the longer 45 mm shank length and to allow the probe to penetrate primate dura. To achieve this, we increased the thickness of the shank by 3.75x, from 24um to 90µm. In addition, since the increased thickness altered the bending profile of the shank, we added stress compensation layers, which help to reduce intrinsic stresses within the material in the shank, and which enabled us to keep the bending within the same range as the rodent probes, despite the 450% increase in length.

For the data reported here, we used the Neuropixels 1.0-NHP probe version with a 125 µm wide, 45 mm long shank, 4416 selectable electrodes or pixels, and a 48 mm^2^ base. To facilitate insertion of the shank into the brain and to minimize dimpling and tissue damage (*1*), the 20° top-plane chisel tapered shanks are mechanically ground to a 25° bevel angle on the side plane using a modified pipette micro grinder (Narishige EG-402). This procedure results in a tip which is sharp along both axes (Fig. 1G), allowing insertion through the dura in many conditions with reduced penetration forces. Additional discussion of insertion mechanics, methods, and hardware are provided in methods and documented in a hardware and methods <u>wiki</u>.

### Scientific applications

The dense, high-channel count, programmable sites of Neuropixels 1.0-NHP provide a number of advantages relative to existing neural recording technologies appropriate for primates and other large animal models in neuroscience. First, the large number of simultaneously-recordable channels represents a transformative capability in itself. Large-scale recordings: A) permit rapid surveys and mapping of recorded brain regions, B) enable analyses which infer the neural state on single trials from large population recordings, C) make it practical to infer functional connectivity using correlational analyses of spike timing, and D) reduce the time and challenge required to perform experiments. Second, the high density of recording sites enables high quality, automated spike sorting (*25*) when recording from one or both columns within a 3.84 mm bank of electrodes (Fig. 1C,H), and high density enables continuous tracking of neurons in the event of drifting motion between the probe and tissue (*14, 15*).

Third, users can programmatically select to record with full density from one column in each of two banks, for 7.68 mm total length of high-quality single unit recordings or programmatically specify alternate layouts (see Supplementary Fig. S1 for site selection rules). Programmable site selection allows experimenters to decouple the process of optimizing a recording location from positioning the probe. This capability allows experimenters to survey neural activity along the entire length of the probe, without moving it in the brain, in order to map the relative position of electrophysiological features. This procedure can reduce or eliminate ambiguity of the probe’s recording depth and location with respect to a target brain area. In addition to enabling large-scale surveys, the programmability also allows experimenters to leave the probe in place to settle prior to an experiment, in order to improve positional stability during the recording and to reduce the impact of any transient tissue response to insertion on subsequent recordings.

We illustrate these collective advantages using example recordings in different macaque brain structures in pursuit of diverse topics, including: 1) retinotopic organization of extrastriate visual cortex, 2) neural dynamics throughout the motor system, 3) face recognition in face patches of inferotemporal (IT) cortex, and 4) neural signals underlying decision making in posterior parietal cortex.

#### Dense recordings throughout primate visual cortex

More than half of the macaque neocortex is visual in function (*26*), and a multitude of visual areas, containing varied retinotopic organization and neurons with distinct feature-selective properties (e.g. motion, color, etc.), lie beyond the primary visual cortex (V1). Most of these visual areas, however, are distributed throughout the brain and located deep within the convolutions of the occipital, temporal and parietal lobes (Fig. 2A). As a consequence of this, and the limitations of prior single-neuron recording technologies, the majority of electrophysiological studies in visual neuroscience have focused on only a subset of visual areas. Fewer than half of the identified visual areas have been well-studied (e.g. areas V4, MT), while most (e.g. DP, V3A, FST, PO) have only been sparsely investigated, which is surprising given the clear similarities between the macaque and human visual systems (*27, 28*). A more thorough and systematic investigation of neural representation across the numerous contributing regions is only practical with technologies that enable large-scale surveys via simultaneous population recordings, with the capability of accessing both superficial and deep structures. Our initial tests with the Neuropixels 1.0-NHP probe demonstrated that it is well-suited for that purpose.

**Figure 2.**
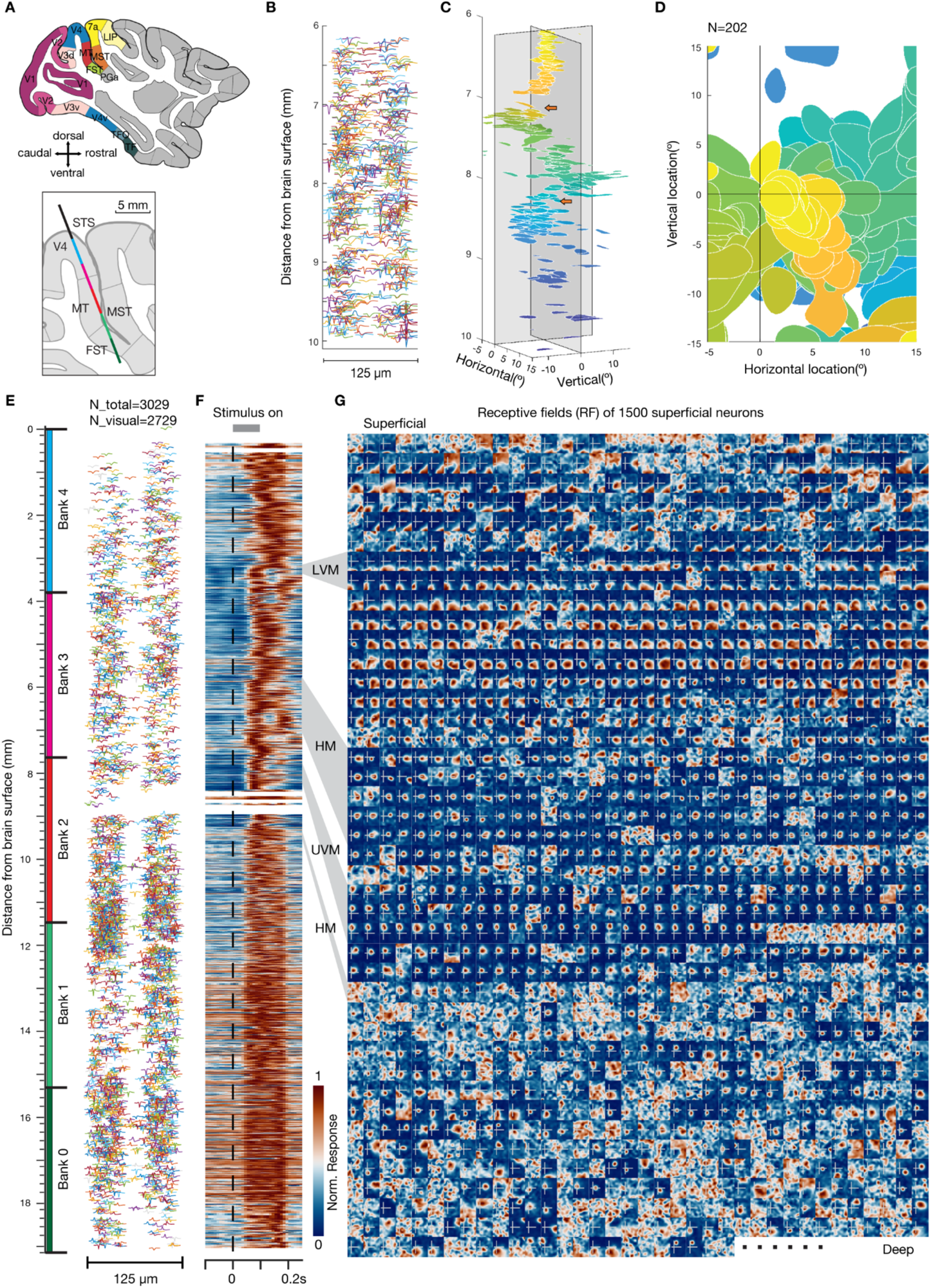
Single and multi-bank recordings across multiple visual cortical areas. **(A)** Visual areas within macaque neocortex shown in a sagittal section. The inset shows the estimated probe trajectory of one multi-bank recording. **(B)** Spike waveforms of single neurons recorded across a single bank of (384) electrode contacts (3.84 mm) shown at their measured location on the probe surface. **(C)** Distribution of receptive fields (RFs) of 202 visually responsive neurons across cortical depth in a single-bank recording. Arrows denote abrupt changes in RF progressions and putative visual area boundaries. **(D)** Top-view of (C) illustrating the coverage of RFs across the contralateral visual field. **(E)** Spike waveforms of 3,029 single neurons recorded across 5 banks of electrode contacts (∼19 mm) shown at their measured location on the probe surface. **(F)** Heat map of stimulus evoked responses for all 2729 visually responsive neurons. Each neuron is plotted at its corresponding cortical depth. Dashed black line denotes stimulus onset; gray line at top denotes (0.1 sec) duration. Gray shading on the right denotes depths where RFs fell on the lower vertical meridian (LVM), horizontal meridian (HM), or upper vertical meridian (UVM). **(G)** RF heat maps for 1500 of the superficial most neurons, indicating visual field locations where stimuli evoked responses for each single neuron. White crosshair in each map denotes the estimated horizontal and vertical meridians. RFs are arranged in a 42 (rows) by 36 (columns) array.

During individual experimental sessions, the activity of thousands of single neurons across multiple visual cortical areas could be recorded using a single NHP probe. Figure 2B shows the spike waveforms of 202 neurons recorded from one bank of electrodes spanning 6–10 mm below the cortical surface. As anticipated, the location of the neurons’ visual receptive fields (RFs) varied as a function of the position along the Neuropixels probe. The RFs shifted in an orderly way for stretches of approximately 1 mm, consistent with a topographic representation, and then shifted abruptly, reflecting likely transitions between different retinotopic visual areas (e.g., refs. (*29–31*); Fig. 2C, Supplementary Fig. S2A). Across the full depth, RFs tiled much of the contralateral hemifield, and included some of ipsilateral visual space as well (Fig. 2D, Supplementary Fig. S2B).

In other sessions, we recorded from up to five probe banks spanning 0–19 mm beneath the pial surface (Fig. 2E). In one example session, 3,029 single neurons were recorded, of which 2,729 neurons were visually responsive, exhibiting short-latency responses after stimulus onset (Fig. 2F). As with the single-bank recordings (Fig. 2C,D), neuronal RFs shifted gradually for contiguous stretches, punctuated by abrupt changes at specific depths. In the example shown (Fig. 2G), RFs at more superficial sites (0-3 mm) were located at more eccentric locations of the visual field, and then abruptly shifted towards the center and closer to the lower vertical meridian (LVM; ∼3 mm). At the same location, neurons became more selective to the direction of motion (Supplementary Fig. S2D), suggesting a transition from area V4 to areas MT or MST. After that, RFs were located more centrally at the lower contralateral visual field and were observed across several mm. At deeper sites (∼6–7 mm), smaller RFs clustered near the horizontal meridian (HM) for more than 1 mm, then quickly shifted toward the upper vertical meridian (UVM; ∼8 mm). Finally, at the deepest sites (>10 mm), RFs generally became larger and much less well defined. These data illustrate how the Neuropixels 1.0-NHP’s dense sampling and single-unit resolution facilitates large-scale and unbiased mapping of the response properties of neurons across multiple visual areas in the primate brain.

#### Large-scale recording throughout the motor system

Next, we demonstrate the utility of this technology for studying the multiple brain areas involved in movement control. Primary motor cortex (M1) is situated at the rostral bank of the central sulcus and extends along the precentral gyrus. Sulcal M1 contains the densest projections of descending corticomotoneuronal cells and corticospinal neurons, which collectively are understood to convey the primary efferent signals from the brain to the periphery (*32–34*). Constraints of existing technology have led to two broad limitations in studies of the motor system. First, motor electrophysiologists have been forced to choose between simultaneous recording from populations of superficial neurons in gyral motor cortex (PMd and rostral M1) using utah arrays, or alternatively, recording fewer neurons in sulcal M1 using single-wire electrodes or passive arrays of 16–32 contacts (e.g., Plexon S-probes or Microprobes Floating Microwire Arrays (*17*)). Recording from large populations of neurons in sulcal M1 has not been feasible.

Second, the motor cortex is only one part of an extensive network of cortical and subcortical structures involved in generating movement (*35*). Many investigations of the motor system focus on the primary motor cortex, and comparably fewer experiments have systematically investigated neural responses from the numerous additional brain structures involved in planning and controlling movements, in part due to the challenge of obtaining large-scale datasets in subcortical structures in primates. Areas such as the supplementary motor area (SMA) and the basal ganglia (BG) are understood to be important for planning and controlling movements, but systematic investigation of the functional roles and interactions between these regions is hampered by the challenge of simultaneously recording from multiple areas.

We developed an insertion system capable of simultaneous insertion of multiple Neuropixels 1.0-NHP probes to superficial and deep structures of rhesus macaques. We tested this approach using a motor behavioral task, in which a monkey used isometric forces to track the position of a scrolling path of dots (Fig. 3A, task described in ref. (*36*)). During this task, we recorded from the primary motor and premotor cortex (M1 and PMd, Fig. 3B) while the monkey generated repeated forces across multiple trials (Fig. 3D). Single neurons exhibit a diversity of temporal dynamics throughout the motor behavior (Fig. 3E–F), and the highest-variance principal components illustrate population dynamics throughout the motor behavior (Fig 3G). Predicting the arm force via linear regression from the neural population reveals improved model performance as more neurons are included, and performance does not saturate when including all recorded neurons (360) for an example session (Fig. 3H). Despite the apparent simplicity of a one-dimensional force tracking task, it is necessary to sample from many hundreds of neurons in order to capture a complete portrait of the population-level neural dynamics.

**Figure 3.**
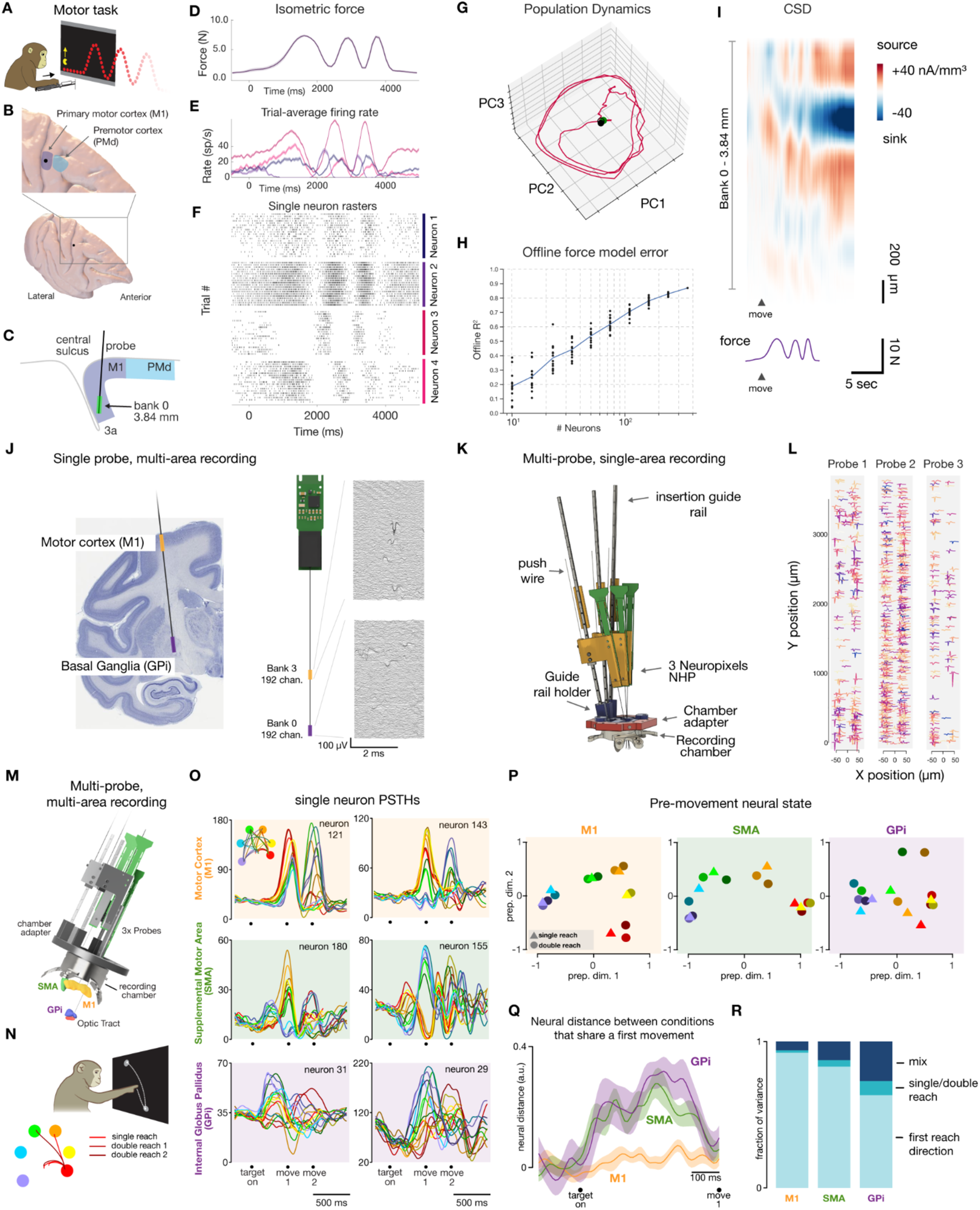
Large scale recording throughout the rhesus motor system. **(A)** Isometric force path-tracking (pacman) behavioral task. **(B)** Probe insertion point in motor cortex visualized on macaque brain rendering. **(C)** Schematic recording target in sulcal M1, sagittal section. **(D)** Force generated by a monkey’s arm during the pacman task. **(E)** Trial-averaged firing rate of four example neurons. **(F)** Single-trial spiking rasters of four example neurons. **(G)** Low-dimensional PCA trajectory of neural activity. **(H)** Linear model force prediction accuracy as a function of the number of neurons included in the analysis. **(I)** Current source density (CSD) calculated using LFP-band signals. **(J)** simultaneous dual region recording using a single probe in M1 and GPi. **(K)** Application 2: multi probe recording within a single brain region. 3D model of insertion setup for three probes with trajectories targeting close-packing arrangement in M1. **(L)** Waveforms from over 673 simultaneously-recorded units in M1. **(M)** 3D model of multi-region recording using multiple probes in GPi, M1, and SMA with recording chamber and insertion hardware. **(N)** Sequential multi-reach task. Monkeys made standard center-out reaches or rapid double reaches to two peripheral targets. During double reach trials, both targets were displayed during the delay period. **(O)** Trial-averaged firing rates from six example neurons. Activity was aligned to target onset, the onset of the first movement, and (when applicable) the onset of the second movement. **(P)** Neural state shortly before (83 ms) movement onset in M1, SMA, and GPi. This was the time, during the delay period, when cross-condition variance was highest in M1. **(Q)** Population neural distance between conditions that share a first movement in each of M1, SMA, and GPi. Distances were calculated within a twenty-dimensional preparatory space (see *Methods*). **(R)** Fraction of variance explained by neural dimensions identified via dPCA for first reach, single vs. double reach, or a mixture of factors in each brain area.

In many experimental situations, it is desirable to be able to precisely localize the probe within the anatomy of the target region. Current source density (CSD) is often used to infer the recording depth using consistent patterns of sources and sinks if the recording trajectory is orthogonal to the cortical lamina (*11, 37, 38*). The high-density of recording sites on the Neuropixels 1.0-NHP allows for averaging of CSD within local regions, enabling high-quality CSD calculations, revealing spatial and temporal structure throughout the force-tracking task (Fig. 3I, Supplementary Fig. S3). This approach may be used to gain confidence in the recording location, and to map the response properties of individual neurons to specific locations within the cortex or deep brain structures. While a majority of prior work has focused on PMd and M1, the programmable site selection enables simultaneous recording from superficial and deep structures using a single probe. To illustrate this capability, we recorded from superficial motor cortex and GPi in the basal ganglia using 192 channels in each target location (Fig. 3J).

The small form factor of the Neuropixels 1.0-NHP probes and headstages allows for dense packing of multiple probes. For applications requiring large numbers of simultaneously recorded neurons from a single area, several probes can be inserted with less than 1 mm spacing between probe shanks by inserting probes along non-parallel trajectories (Fig. 3K). Figure 3L illustrates recordings from three probes in gyral PMd, yielding 673 neurons, and up to 819 neurons using six probes in a separate session (not shown). This approach is also well suited for simultaneous recording from different brain areas. Figure 3M illustrates a setup used to simultaneously record from M1, the internal globus pallidus (GPi) of the basal ganglia, and the supplementary motor area (SMA) (Fig. 3N). Neurons in all three regions displayed modulated trial-averaged activity patterns during a sequential multi-target reaching task (Fig. 3O). Neural preparatory activity prior to single and double reaches was more similar in M1 for reaches with the same initial movement segment than in SMA or GPi (Fig. 3P). This divergence emerged after targets were presented on screen and persisted until just before the movement began (Fig. 3Q). Analysis of population variance using dPCA reveals that in all three areas, delay-period activity most strongly corresponds to the direction of the first reach. First-reach related variance accounted for 92, 83, and 63% of the total variance captured by dPCA in M1, SMA, and GPi, respectively. Activity in SMA and GPi was also influenced by the identity of the second reach in a two-reach sequence; 17% and 37% of the variance explained by dPCA was represented by dimensions related purely to condition context (single or double reach) or a mixture of first reach identity and context. By comparison, these dimensions did not account for a large fraction of variance in M1 activity (8% of the total variance explained by dPCA). Even this small fraction of neural variance is likely due to kinematic differences between single reaches and the first half of double reaches.

Conveniently recording from very large populations of neurons is itself a transformative capability, as this enables experimenters to perform experiments that may not have been feasible to collect otherwise. Such experiments could either be high-risk, high-reward projects, or simply additional control experiments. This is particularly true for a range of deeper brain regions known to be important for motor control, but which have historically received less attention due to the relative challenge of accessing populations of neurons in these areas. The capabilities enabled by the Neuropixels 1.0-NHP probe extend beyond simply reducing the challenge of obtaining neural data. Simultaneous large-scale population recordings from multiple structures enable identification of the communication between areas, for example via communication subspaces (*40, 41*).

#### Face recognition in IT cortex

Next, we demonstrate use of the probes in the inferotemporal (IT) cortex, a brain region that is challenging to access due to its depth, spanning the lower gyrus and ventral surface of the temporal lobe. IT cortex is a critical brain region supporting high-level object recognition, and has been shown to harbor several discrete networks (*42*), each specialized for a specific class of objects. The network that was discovered first and has been most well-studied in nonhuman primates is the face patch system. This system consists of six discrete patches in each hemisphere (*43*), which are anatomically and functionally connected. Each patch contains a large concentration of cells that respond more strongly to images of faces than to images of other objects. Studying the face patch system has yielded many insights that have transferred to other networks in IT cortex, including increasing view invariance going from posterior to anterior patches and a simple, linear encoding scheme (*42*). As such, this system represents an approachable model for studying high-level object recognition (*44*). The code for facial identity in these patches is understood well enough that images of presented faces can be accurately reconstructed from neural activity of just a few hundred neurons (*45*).

A major remaining puzzle is how different nodes of the face patch hierarchy interact to generate object percepts. To answer this question, it is imperative to record from large populations of neurons in multiple face patches simultaneously to observe the varying dynamics of face patch interactions on a single-trial basis. This is essential since the same image can often invoke different object percepts on different trials (*46*). Here, we recorded with one probe in each of two face patches, middle lateral (ML) and anterior fundus (AF) simultaneously (Fig. 4A). The Neuropixels 1.0-NHP probes recorded responses of 1,127 single- and multi-units across both face patches during a single session (Fig. 4A, right). Changing the visual stimulus to a monkey face yielded a clear visual response across both face patch populations. We measured responses of visually responsive cells to 96 different stimuli containing faces and non-face objects (Fig. 4B–C). A majority of cells in the two patches showed clear face-selectivity. With single-wire tungsten electrodes, this dataset would take about two years to collect, but is now possible in a single two-hour experimental session (*45*). In addition to the gain in efficiency created by this technology, simultaneous recordings of multiple cells and multiple areas allows for investigation of how populations encode object identity in case of uncertain or ambiguous stimuli, where the interpretation of the stimulus may vary from trial to trial but is nevertheless highly coherent on each trial. The anatomical depth of face patches puts them far out of reach for shorter high-density probes. For example, face patch AM sits roughly ∼42 mm from the craniotomy (Fig. 4D) along a conventional insertion trajectory.

**Figure 4.**
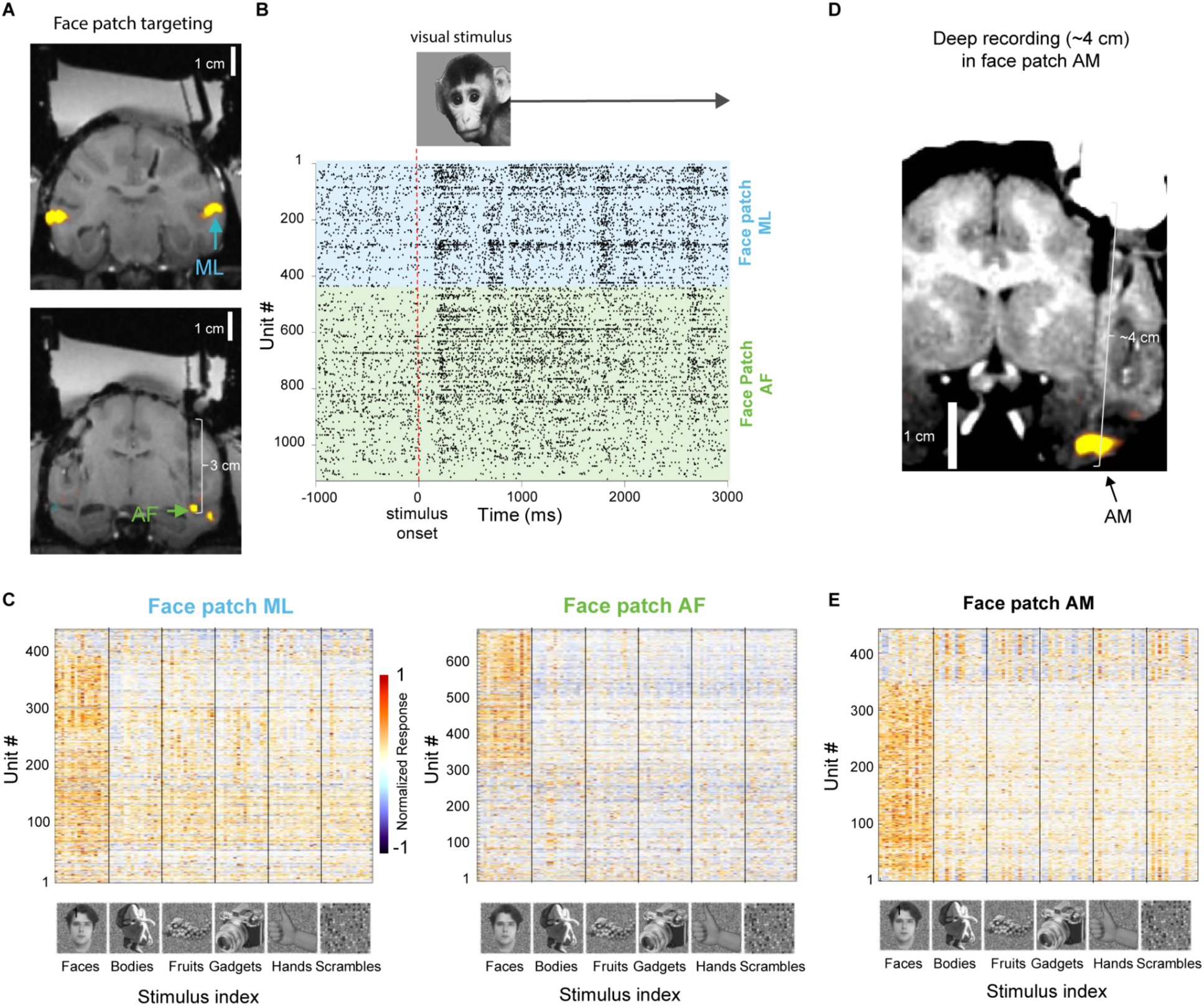
Deep, simultaneous recordings from two face patches in IT cortex. **(A)** Left: Simultaneous targeting of two face patches. Coronal slices from magnetic resonance imaging (MRI) scan show inserted tungsten electrodes used to verify targeting accuracy for subsequent recordings using Neuropixels 1.0-NHP (top: face patch ML, bottom: face patch AF). Color overlays (yellow) illustrate functional magnetic resonance imaging contrast in response to faces vs. objects. **(B)** Response raster for a single stimulus presentation of simultaneously recorded neurons in ML and AF to a monkey face, presented at t=0. Each line in the raster corresponds to a spike from a single neuron or multi-unit cluster (including both well isolated single units and multi-unit clusters. **(C)** Neuropixels 1.0-NHP enables recordings from many face cells simultaneously. These plots show average responses (baseline-subtracted and normalized) of visually responsive cells (rows) to 96 stimuli (columns) from six categories, including faces and other objects. Bottom panel shows exemplar stimuli from each category. The plots include 438 cells or multiunit clusters in ML (left) and 689 in AF (right), out of which a large proportion responds selectively to faces. Units are sorted by channel, revealing that face cells are spatially clustered across the probe. **(D)** Coronal slice of MRI scan of Neuropixels probe targeting the deepest IT face patch AM. The thick shadow is a cannula that was inserted through the dura into the brain. The thin shadow corresponds to the Neuropixels trajectory. **(E)** Same as **C**, for face patch AM (recording performed in a different session from data in **C**).

#### Single-trial correlates of decision making in LIP

For many cognitive functions, the processes that give rise to behavior vary across repetitions of a task. For example, certain perceptual decisions are thought to arise through the accumulation of noisy evidence to a stopping criterion, such that their evolution is unique on each trial. This process is widely observed and known as drift-diffusion (*47, 48*). Its neural correlate has been observed in the lateral intraparietal area (LIP). Neurons in LIP have spatial response fields (RFs; Fig. 5C, top) and represent the accumulated evidence for directing the gaze toward the response field. Limitations in recording technology prevented previous studies from recording many LIP neurons simultaneously, requiring that neural activity be averaged across similar decisions. Such averaging highlights the shared features of activity across decisions (e.g., the “drift” component) but discards the unique dynamics that give rise to each individual decision, specifically the diffusion component.

**Figure 5.**
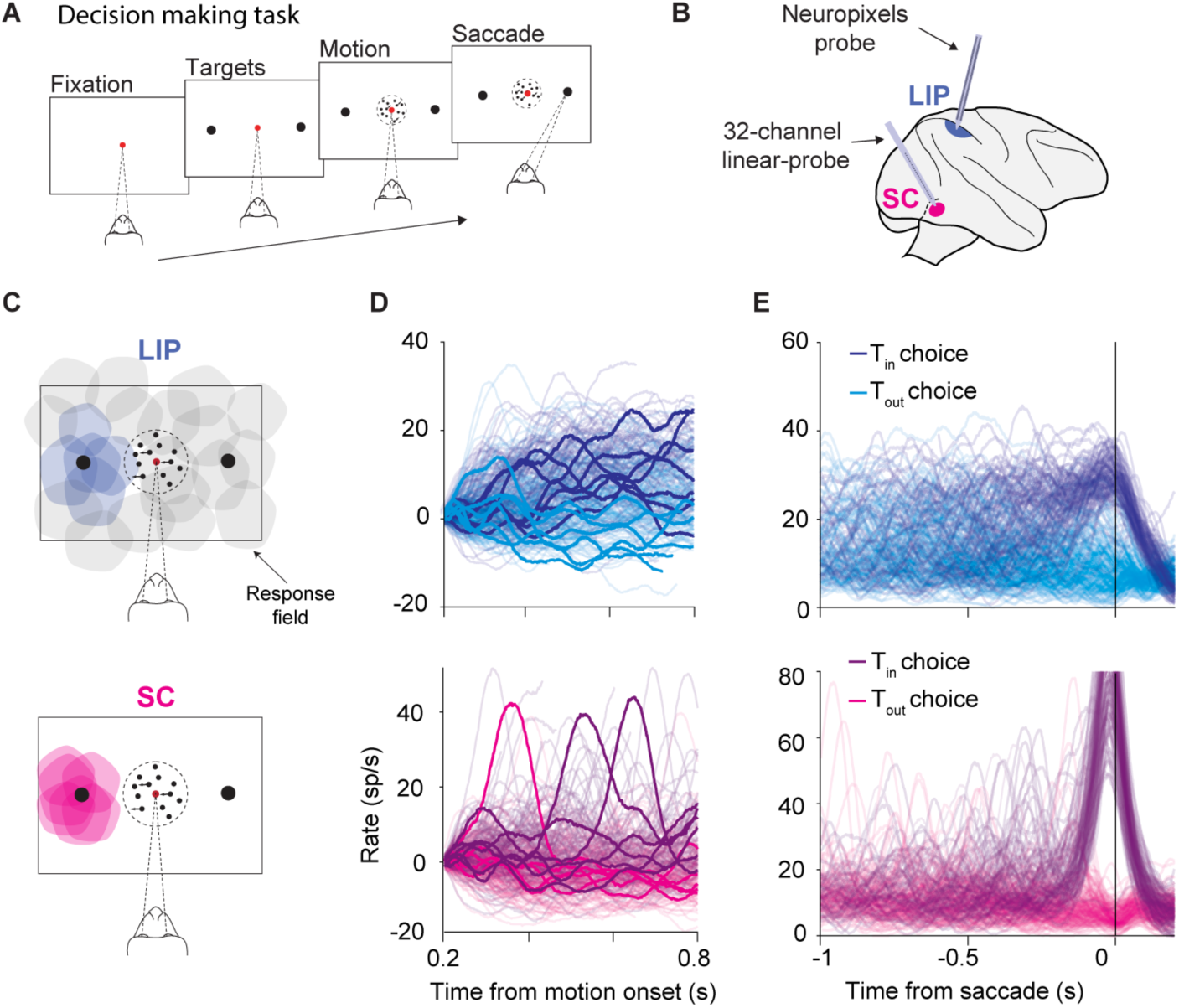
Single-trial dynamics of a decision process in multiple brain regions. **(A)** Task. The monkey must decide the net direction of dynamic random dot motion and indicate its decision by making an saccadic eye movement, whenever ready, from the central fixation point (red) to a left-choice or right-choice target (black). The choice and response times are explained by the accumulation of noisy evidence to criterion level (Gold & Shadlen, 2007). **(B)** Simultaneous recordings in LIP and SC. Populations of neurons were recorded in LIP with a Neuropixels 1.0-NHP probe and in SC (deeper layers) with a multi-channel V-probe (Plexon). **(C)** Response fields (RF) are identified, *post hoc*, from control blocks in which the monkey performed an oculomotor delayed response task. A small fraction of the LIP neurons (top) have RFs that overlap the left-choice target. The SC (*bottom*) has a topographic map, so many neurons have overlapping RFs. The large sample size in LIP facilitates identification of neurons that respond to the same choice target in LIP and SC. **(D)** Single trial activity in LIP (top, n = 17) and SC (bottom, n = 15). Each trace depicts the smoothed firing rate average (Gaussian kernel, σ=25 ms) of the 17 neurons that overlap the left-choice target (Tin) on a single trial, aligned to the onset of the motion stimulus. Rates are offset by the mean firing rate 0.18-0.2 s after motion onset to force traces to begin at zero. The line color indicates the decision for the left-choice target or the right-choice target (Tout) on that trial. A few representative traces are highlighted for clarity **(E)** The same trials as in d aligned to saccade initiation, without baseline offset. Single trial firing rates approximate drift-diffusion in LIP—the accumulation of noisy evidence, whereas single trial firing rates in SC exhibit a large saccadic burst at time of the saccade, preceded by occasional non-saccadic bursts.

Neuropixels recording in LIP reveals the neural correlates of a single decision. These recordings yield 50–250 simultaneously recorded neurons, of which 10–35 share a RF that overlaps one of the contralateral choice targets used by the monkey to report its decision in a dynamic motion discrimination task (Fig. 5A). The average activity of these target-in-RF (T_in_) neurons on a single trial tracks the monkey’s accumulated evidence as it contemplates its options. The signal explains much of the variability in the monkey’s choices and reaction times (*49*) and conforms to drift-diffusion dynamics (Fig. 5D–E, top).

Neuropixels technology also enables multi-area recordings from ensembles of neurons that share common features. For example, neurons in the deeper layers of the superior colliculus (SC) receive input from LIP and, like LIP neurons, also have spatial RFs and decision-related activity. An ideal experiment to understand how the two areas interact is to record simultaneously from populations of neurons in LIP and SC that share the same RF. This experiment is nearly impossible with previous recording technology because of the anatomical organization in LIP and SC. SC is retinotopically organized such that nearby neurons share the same response field, but such organization is absent in LIP. Therefore, it is improbable that a given LIP neuron will have a response field that overlaps with those of a cluster of SC neurons. With patience, one can find one LIP neuron with the desired RF, but the likelihood of encountering more than two is vanishingly small. This challenge is overcome by the large number of neurons yielded by Neuropixels 1.0-NHP probe recording, allowing for *post hoc* identification of neurons with overlapping RFs (Fig. 5C). In the experiment depicted in Figure 5, the response fields of 17 of the 203 LIP neurons overlap the left-choice target as well as 15 simultaneously recorded neurons in the SC. Unlike LIP, single trial analysis of the SC population reveals dynamics that are not consistent with drift-diffusion (Figure 5d-e, bottom). Instead, SC exhibits bursting dynamics, which were found to be related to the implementation of a threshold computation (*50*). The distinct dynamics in LIP and SC during decision making were only observable through single-trial analyses, the resolution of which are greatly improved with the high yield of the Neuropixels 1.0-NHP recording.

#### Inferring functional connectivity with high density recordings

Understanding how the anatomical structure of specific neural circuits implements neural computations remains an important but elusive goal of systems neuroscience. One step towards connecting disparate levels of experimental inquiry is mapping functional connections, or inferred connectivity between neurons using correlative measures of spike timing (*6, 11, 51*). This is often impractical or extremely challenging when only recording from a small number of neurons, since the likelihood of recording from a connected pair of neurons can be quite low. The probability of recording from connected pairs of neurons depends on a number of factors, including the details of anatomy in a specific species and brain region. Broadly speaking, however, the probability of successfully recording two functionally-connected neurons increases with the square of the total number of neurons recorded.

The Neuropixels 1.0-NHP probe typically yields 200–450 (and sometimes more) neurons when recording with 384 channels in cortical tissue. Applying the same methodology established in ref. (*6*) to 13 sessions from rhesus PMd and M1 yielded 111 ± 89 putative connected pairs per session, and a connection probability of 0.73% ± 0.61%. Fig. 6A shows three example jitter-corrected cross correlogram plots between pairs of neurons with significant peaks in the CCG. In many examples the CCG peak is lagged between one neuron relative to the other, consistent with a 1–2 ms synaptic delay. For other neuron pairs, the CCG peak is synchronous between the two neurons, suggesting that they may receive common input (e.g., Fig 6a, top right panel). Supplementary Fig. 4 illustrates the distribution of spike timing delays for 479 neurons from an example recording session. Nearby neurons are more likely to exhibit functional connections than neurons located further apart. Using this approach, we can map the full set of putative connections for a given recording across cortical lamina (Fig. 6B).

**Figure 6.**
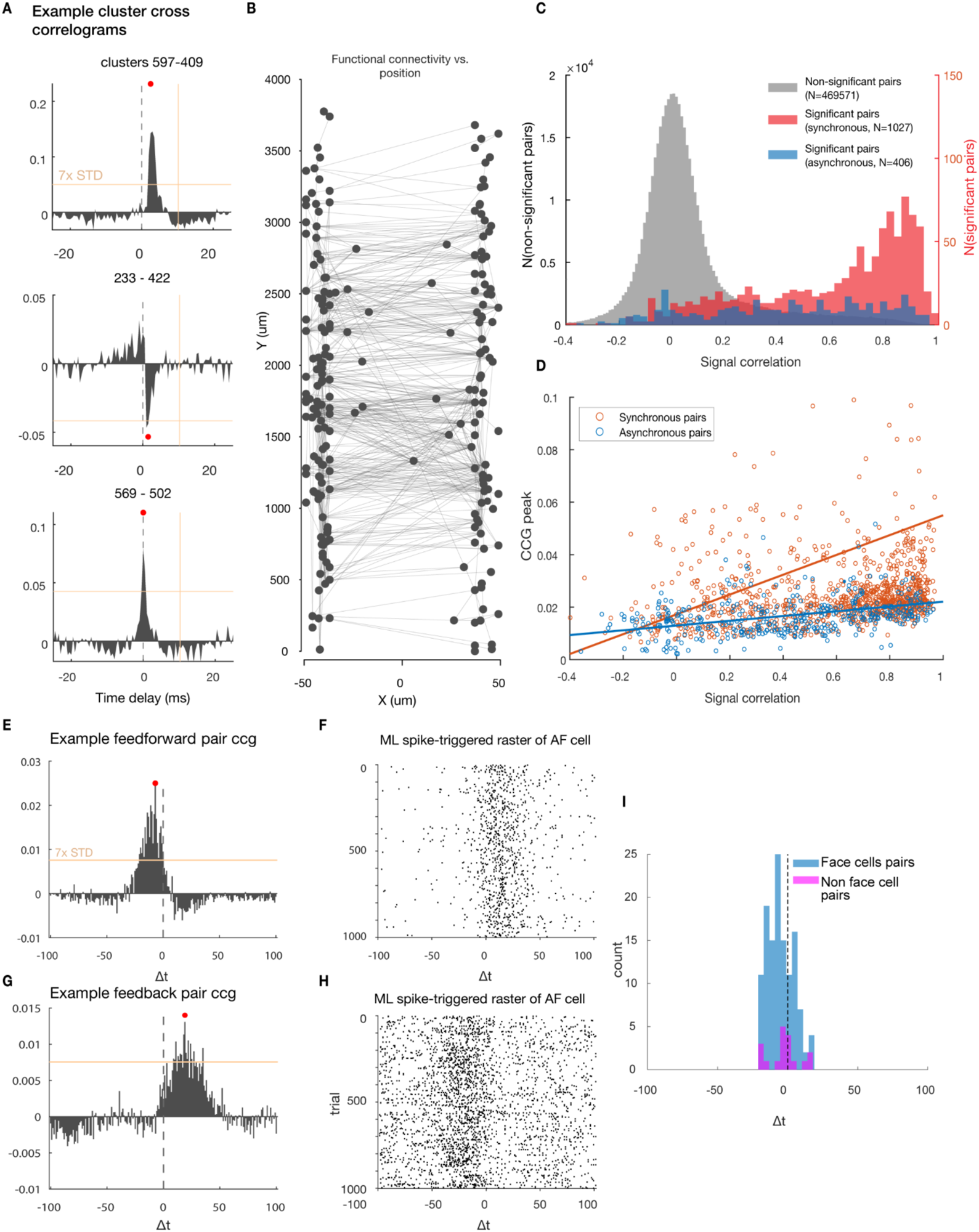
Inferring functional connectivity from unit cross-correlation and high-density recording. **(A)** Jitter-corrected cross correlograms of four example pairs of neurons exhibiting significant correlations in spike timing. **(B)** Diagram of functionally connected neuron pairs from one example session, with neurons ordered by depth along the probe. **(C)** Distribution of signal correlation for pairs of neurons with different CCG types in the visual cortex. **(D)** relationship between signal correlation and peak value of CCG. **(E-F),** Example putative connected cell pair identified using two probes in area ML and AF with putative feedforward connection. **(G-H),** Example putative connected cell pair identified using two probes in area ML and AF with putative feedback connection. **(I),** The population of functionally connected cells between AL and MF regions is dominated by cells that respond to faces.

In addition to the cortical distance, we further assessed the dependence of functional connectivity on the tuning similarity between neuronal pairs for data recorded in the visual cortex. For example, neuron populations with diverse receptive fields (RFs) obtained from multiple visual areas (Fig. 2) allow us to quantitatively determine the extent to which functional connections between neurons depend on their RFs overlapping. Using signal correlation (r_sig_) as a measure of such tuning similarity in visual fields (*52*), we show that functionally connected neuron pairs exhibit higher r_sig_ compared with non-significant pairs (Fig. 6C). Specifically, synchronous pairs (putatively receiving common inputs) tend to share highly overlapping RFs (r_sig_ mean=0.60, p<10^-4^ compared with non-significant pairs), while asynchronous pairs (putatively exhibiting synaptical connections) more likely to share moderately overlapping RFs (r_sig_ mean=0.43, p<10^-4^ compared with non-significant pairs, and p<10^-4^ compared with synchronous pairs). Moreover, the amplitude of the significant CCG peaks is positively correlated with the r_sig_ (Fig. 6D), indicating that neurons with similar RFs tend to have stronger functional connections, which is consistent with previous studies (e.g., (*53, 54*)).

This same methodology can be applied to assess functional connectivity between multiple simultaneously recorded regions recording using separate probes to assess feedforward and feedback connectivity between two regions. Fig. 6E shows the jitter-corrected CCG for a putatively connected pair of neurons where one neuron is located in face patch ML and the other in face patch AF in IT cortex. Fig 6F shows a single-trial spike raster of the putative post-synaptic neuron in area AF, triggered on spikes of the neuron in ML, illustrating a putative feedforward connection, while Fig 6H illustrates a putative feedback connection with opposite timing response from the cell pair shown in Fig. 6G. Remarkably, pairs of face cells were over 10 times as likely to be connected (1.6%) as other pairs of non-face responsive cells (0.13%) between ML and AF (Fig. 6I).

## Discussion

We have presented a new recording technology and suite of techniques to enable electrophysiological recordings using high-density integrated silicon electrodes in rhesus and other nonhuman primates. This technology enables large-scale recordings from populations of hundreds of neurons from deep structures in brain areas that are inaccessible using alternate technologies. The key methodological advance in the Neuropixels 1.0-NHP probe is the longer recording shank, which required developing techniques to adapt photolithographic silicon manufacturing methods to allow for “stitching” across multiple reticles. This advance allows for the manufacturing of monolithic silicon devices that span across multiple reticles to achieve larger sizes than could otherwise be manufactured. Creating a long and thin probe shank also required developing approaches for reducing bending due to internal stresses within the shank.

This technology combines the advantages of multiple approaches - recording with single-neuron spatial resolution and single-spike temporal resolution, while providing recording access to the majority of the macaque brain. Programmable site selection enables recording from multiple brain structures using a single probe, as well as surveying multiple recording sites along the shank without moving the probe. The combination of the compact form factor, commercially-available and turn-key recording hardware, and comparably modest cost enable straightforward scaling in the size of simultaneously recorded neural populations. These capabilities may be essential for achieving accurate estimates of neural dynamics on single trials, or for estimating the value of small-variance neural signals embedded in the neural population response.

The high spatial resolution offers a number of advantages over sparser sampling, including high quality single-unit isolation, automated drift correction (e.g., refs. (*25, 55*), Supplementary Fig. S5), and localizing the position and depth of the recording electrodes within a brain structure (e.g., inferring probe depth with respect to cortical lamina) using current source density or other features of the recording. The high density also offers additional advantages not described here, such as identifying putative neuron subclasses using extracellular waveforms (*5, 14, 56*).

The probe is now commercially available and integrates seamlessly with the existing set of community-supported hardware and software tools for Neuropixels probes. The Neuropixels 1.0-NHP recording system is straightforward to set up and to integrate with other experimental hardware, like behavior or stimulus control computers. Combining the low total system cost (roughly $7–15k) and large-scale recordings enables a dramatic reduction in the recording cost per neuron acquired relative to existing technologies.

While highly capable, the Neuropixels 1.0-NHP probes are limited in several ways. First, this technology is not optimized for simultaneous, dense sampling across a wide swath of cortex. For applications requiring horizontal sampling, planar recording technologies like Utah arrays or two-photon calcium imaging may be more appropriate. Second, in contrast with many passive electrodes, it is not currently possible to use the Neuropixels probe to deliver intracortical microstimulation (ICMS), though future versions of the probe may add this functionality. The Neuropixels technology is, however, capable of recording while stimulating through external electrodes, as recently demonstrated by O’Shea, Duncker, et al. (*13*). Third, the Neuropixels 1.0-NHP design is not explicitly optimized for chronic implantation. While it is likely possible to leave the probe in place over multiple days or sessions, this possible capability remains untested and requires novel implant designs. The probe base contains active electronics, and is not designed for implantation under dura. As such, a chronic implant design may require mounting the probe in a manner that allows it to mechanically “float” with the brain, to prevent relative motion between the probe and the tissue as the brain moves. As such, the current probe is most appropriate for acute recordings, though it conceivably could be implanted for sub-chronic (multiple-week) recordings with appropriate insertion methods and hardware. Lastly, we note that while it is theoretically possible to insert the entire 45 mm long shank into the brain, inserting a probe this deep introduces additional practical challenges to overcome—primarily a requirement for precise alignment of the probe’s insertion axis with the insertion location. A thorough discussion of these considerations is presented in a user’s wiki https://github.com/etrautmann/Neuropixels-NHP-hardware/wiki.

Taken together, these methodological advances enable new classes of neuroscientific experiments in large animal models and provide a viable scaling path towards recording throughout the whole brain.

## Acknowledgements

We thank the support of the Howard Hughes Medical Institute, who funded the development of the probe. We thank Yanina Pavlova, Danielle Abreu Lopes, Stephen Cital, Cornel Duhaney, Brian Madeira, Mackenzie Risch, and Michelle Wechsler for surgical assistance and expert veterinary care. for their assistance in the planning and execution of surgeries, animal training and general support, and Stephen Ryu for surgical expertise. We thank Bob Schneeveis and Tanya Tabachnik for engineering assistance. In addition, we thank Columbia University’s ICM for the quality of care they provide for our animals, especially during the pandemic and lockdown. We thank Wei-lung Sun, for probe testing and software development (HHMI Janelia).

## Author Contributions

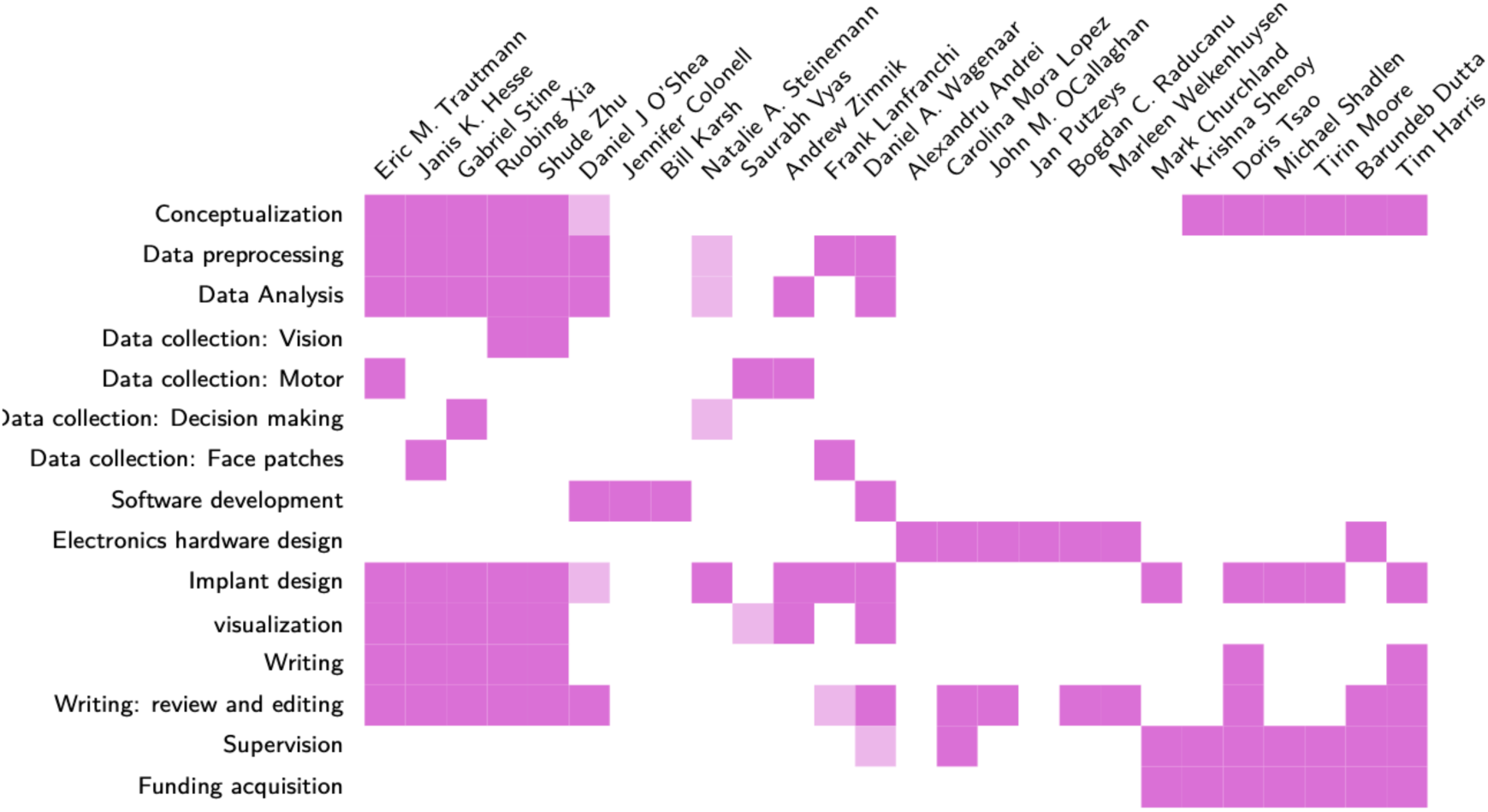

## Data and materials availability

Data and code to replicate key analyses will be made available upon publication.

## Materials and Methods Summary

### Probe design and recording system

The Neuropixels 1.0-NHP probe consists of an integrated base and “shank” fabricated as a monolithic piece of silicon using a 130 nm CMOS lithography process. The 6 mm x 9 mm base is mounted to a 7.2 x 23 mm PCB, which is attached to a 7.2 x 40 mm long flexible PCB. This flex PCB plugs into a ZIF connector on a headstage (15 x 16 mm, 900 mg), which is connected to a PXIe controller mounted in an PXIe chassis using a 5 m twisted-wire cable. The base electronics, headstage, cable, PXIe system, and software are identical to the Neuropixels 1.0 probe. Data collection was performed using SpikeGLX software (*57*) and the system is fully compatible with OpenEphys software.

The Neuropixels 1.0-NHP probe is manufactured in two variants: 1) 45 mm long x 125 µm wide x 90 µm thick, featuring 4416 electrodes comprising 11.5 banks of 384 channels each; and 2) 25 mm long, 125 µm wide, and 60 µm thick, featuring 2496 electrodes comprising 6.5 banks of 384 channels. We note here that the commercial release of these probes will feature electrodes distributed in two aligned vertical columns, in contrast to the “zig-zag” columns described in this manuscript, in order to optimize data collection for automated drift correction to enhance automated spike sorting. We also note that a third variant, identical to Neuropixels 1.0 but with a thicker shank (100 µm vs. 25 µm), is also planned for commercial release.

Recording sites are 12 µm x 12 µm, made of titanium nitride, and have an impedance of 150 kΩ at 1kHz. The tips of the probes were mechanically beveled to a 25° angle using the Narishige EG-402 micropipette beveler. During recordings, electrical measurements were referenced to either: 1) the large electrical reference point on the tip of the electrode, 2) an external electrical reference wire placed within the recording chamber, or 3) a stainless steel guide tube cannula. Electrical signals are digitized and recorded separately for the action potential (AP) band (10 bits, 30 kHz, 5.7 µV mean input-referred noise) and local field potential (LFP) band (10 bits, 2.5 kHz).

Recording sites are programmatically selectable with some constraints on site-selection. See Supplementary Fig. S1 for a description of site selection rules and common configurations. Spike sorting was performed using Kilosort 2.5 and Kilosort 3.0, and results were curated using Phy. Analysis was performed using custom scripts written in Matlab and Python, leveraging the open source software package neuropixels-utils: (https://github.com/djoshea/neuropixel-utils).

### Probe insertion

Several distinct methods were used to mount and insert probes, guided by the unique constraints of inserting probes to different depths, and depending on the recording chambers and mechanical access available for different primates used in these studies, as well as the existing hardware used by each of four distinct research groups. For single probe insertions, probes were mounted using custom adapters to a commercially available electrode drive (e.g., Narishige corp.), and inserted through a blunt guide tube for superficial recordings, and a sharp penetrating guide tube for deeper recordings. When using a non-penetrating guide tube, the dura was typically penetrated with a tungsten electrode prior to using a Neuropixels probe to create a small perforation in the dura to ease insertion. When inserting electrodes to deep targets (>20 mm), the alignment between the drive axis and the probe shank is essential for enabling safe insertion, as misalignment can cause the probe to break. For this application, we developed several approaches to maintain precise alignment of the probe and drive axis. First, we employed a linear rail bearing (IKO International) and custom 3D printed fixture to maintain precise alignment of the insertion trajectory. This approach is discussed in detail in the accompanying Neuropixels-NHP wiki (https://github.com/cortex-lab/neuropixels/wiki).

For the experiments shown in figure 4, we developed a dovetail rail system that maintains precise alignment between a penetrating guide tube and the Neuropixels probe. The choice of appropriate insertion method depends on the mechanical constraints introduced by the recording chamber design, the depth of recording targets, number of simultaneous probes required, and choice of penetrating or non-penetrating guide tube. The interaction of these constraints and a more thorough discussion of insertion approaches is provided on the Neuropixels users wiki. Open-source designs for mechanical mounting components for Neuropixels-1.0-NHP to drives from Narishige, NAN, and other systems are available in a public repository: https://github.com/etrautmann/Neuropixels-NHP-hardware.

### Visual cortex recordings and analysis

Two male adult rhesus monkeys (*Macaca Mulatta*, 11 and 16 kg), monkey T and monkey H, served as experimental subjects. Each animal was surgically implanted with a titanium head post, and a cylindrical titanium recording chamber (30 mm diameter). In each animal, the placement of the recording chamber was centered at ∼17 mm from the midline and ∼7 mm behind ear-bar-zero, and craniotomy was performed, allowing access to multiple visual areas in the superior temporal sulcus (STS). All surgeries were conducted using aseptic techniques under general anesthesia and analgesics were provided during post-surgical recovery.

We measured visual RFs by randomly presenting a single-probe stimulus out of either a 7 (H) * 11 (V) probe grid extending 18 (H) * 30 (V) degree of visual angles (dva)(monkey T), or a 14 (H) *17 (V) probe grid extending 26 (H) * 32 (V) dva (monkey H). The probes consisted of a circular drifting Gabor gratings (2 deg in diameter, 0.5 cycle/deg in spatial frequency, 4 deg/sec in speed, 100% Michelson contrast) and was presented for a duration of 0.1 sec. Monkeys were recorded with a drop of juice if they maintained fixating at the fixation spot throughout the trial.

For a given probe location, we obtained the neuronal activity by counting all the spikes during the stimulus presentation period, accounted for by a time delay of 50 ms. Neuronal RFs were defined as the probe locations that elicited more than 90% of the peak visual responses.

### Motor cortex recordings and analysis

Details of the pacman behavioral task and experimental hardware presented in Fig. 3 are described in (*58*). Three monkeys (*Macaca Mulatta)* served as experimental subjects. In each, a head post and recording chamber were implanted over premotor and primary motor cortex using aseptic surgical procedures and general anesthesia. Placement of the chambers were guided using structural MRI. Recordings in Monkey C were performed using a standard 19 mm plastic recording chamber (Christ Inc.), while Monkey I and J were implanted with custom low-profile footed titanium chambers (Rogue Research).

We conducted 13 sessions in Monkey C, and 10 sessions in Monkey I, targeting sulcal and gyral M1 and PMd. On a subset of 15 sessions in Monkey C, we also targeted GPi in the basal ganglia. In Monkey J, we report data from one session while simultaneously recording in GPi, SMA, and M1.

For monkey C, the Neuropixels 1.0-NHP probe was held using a standard 0.25” dovetail mount rod with a custom adapter to mount it to a hydraulic drive (Narishige Inc.). A 21 gauge blunt guide tube, 25 mm in length, was held using a custom fixture and placed over the desired recording location. The dura was then penetrated with a tungsten electrode (FHC, size E), which was bent at 27 mm to prevent the tip from inserting further than 2 mm past the end of the guide tube. This electrode was inserted manually via forceps, once or several times, as necessary, which also provided feedback on the depth and difficulty of penetrating the dura. The Neuropixels 1.0-NHP probe was then aligned using the Narishige tower XY stage, lowered into the guide tube, and carefully monitored to ensure that the tip of the probe was aligned with the dural penetration. This procedure sometimes took several attempts to find the correct insertion point, but generally was successful in less than a few minutes.

For Monkeys I and J, the Neuropixels 1.0-NHP probe was held using a custom fixture mounted to a linear rail bearing (IKO Inc.). This apparatus was designed to enable close packing of many probes, and to solve the challenge of precisely targeting structures deep in the brain without trial and error. The linear rail is mounted in a custom 3D printed base, which mounts directly to the recording chamber. The geometry of the 3D printed base component determines the insertion trajectories and prevents mechanical interference between probes and the chamber. This base also provides support for either sharp or blunt guide tubes, as required. In general, blunt guide tubes were preferred, but if necessary sharp guide tubes were sometimes used when the dura had become thicker and difficult to penetrate. The linear bearing was connected to a commercial drive system (NAN Inc.) via a ∼50 mm long, .508 mm stainless steel wire, which provided rigid connection between the Nan drive electrode mount and the Neuropixels probe mounted on the rail bearing, while allowing a small amount of misalignment between the drive axis and the insertion axis. This apparatus greatly simplifies the procedure of using many probes in a small space, while not relying on commercial drives to provide the mechanical rigidity required to safely insert a delicate probe. Additional details on the custom hardware is provided in the <u>Neuropixels 1.0-NHP user wiki</u>.

Spike sorting was performed using Kilosort 2.5 and manually curated using Phy. PCA trajectories were calculated after smoothing spikes with a 25 ms gaussian kernel and averaging across successful trials. Offline force model prediction performance was computed using a 50 ms time lag between arm force and neural activity. Neurons were randomly sub-selected, and 80% of trials from six target conditions were used to train a linear regression model in Python using scikit-learn, while the remaining 20% of trials were used to calculate model performance. Ten iterations were performed for each level of neurons retained. Details of the current source density analysis are presented further below. The dPCA analysis was run on a [# Neurons x reach direction x context (single/double reach)] 3D tensor. Each entry in that tensor was the trial averaged firing rate of a neuron averaged over a 10 ms window centered on a time 83 ms before movement onset, when preparatory activity in M1 was the strongest. The fraction of variance explained used 10 dPCA dimensions, which accounted for 99%, 99%, and 98% of the total neural variance during this window of time in M1, SMA, and GPi respectively.

### LIP recordings and analysis

Details of the collection and analysis of the data presented in Figure 5 are described in(*50*). Two monkeys (*Macaca Mulatta*, 8–11 kg) served as experimental subjects. In each, a head post and two recording chambers were implanted using aseptic surgical procedures and general anesthesia. Placement of the LIP chamber was guided by structural MRI. The SC chamber was placed on the midline and angled back 38° from vertical in the anterior-posterior axis.

We conducted eight recording sessions in which activity in LIP and SC was recorded simultaneously. In LIP we used a single Neuropixels 1.0-NHP probe, yielding 54–203 single units per session. In SC, we used 16-, 24-, and 32-channel V-probes (Plexon) with 50–100 µm electrode spacing, yielding 13–36 single units per session. In each session, we first lowered the SC probe and approximated the response fields (RFs) of SC neurons using a few dozen trials of a delayed saccade task. Because our penetrations were approximately normal to the retinotopic map in SC, the RFs of the SC neurons were highly similar within a session. We proceeded only if the center of the RFs were at least 7° eccentric in order to ensure minimal overlap with the motion stimulus.

If the RF locations in SC were suitable, we then lowered the Neuropixels probe into LIP through a dura-penetrating, stainless steel guide tube (23G) at 5 µm/s using a MEM microdrive (Thomas Recording) that was attached to a chamber-mounted, three-axis micromanipulator. Custom-designed adapters were used for mounting the Neuropixels probe onto the drive (see <u>wiki</u>). Once the target depth was reached (∼10 mm below the dura), we allowed for 15–30 minutes of settling time to facilitate recording stability. In order to precisely measure RF locations in both areas, the monkeys performed 100–500 trials of the delayed saccade task and LIP neurons with RFs that overlapped those of the SC neurons were identified post-hoc. Finally, the monkeys performed a reaction-time RDM motion discrimination task until satiated (typically 1,500–3,000 trials).

Neurons in both areas were sorted using Kilosort 2.0 and manually curated in Phy. We restricted our analysis of the LIP data to neurons with RFs that overlapped those of the simultaneously recorded SC neurons. (164 of 1,084 total LIP neurons). Spike trains were discretized into 1 ms bins and convolved with a Gaussian kernel (σ = 25 ms) to produce the single-trial activity traces depicted in Figure 5d and e.

### Face patch recordings and analysis

Three monkeys (*macaca mulatta*) served as experimental subjects. Each animal was surgically implanted with an MRI-compatible Ultem head post, and a large rectangular recording chamber (61 x 46 mm diameter, 65 x 50 mm diameter, and 58 x 61 mm diameter respectively), covering most of the animal’s acrylic implant. Monkeys were trained to passively fixate on a spot for juice reward while visual stimuli of 5° size, such as images of faces or objects, were presented on an LCD screen (Acer). We targeted face patches ML and AF (monkey 1), face patches MF and AL (monkey 2), and face patches MF and AM (monkey 3) in the IT cortex for electrophysiological recordings. Face patches were identified using fMRI. Monkeys were scanned in a 3T scanner (Siemens), as described previously (*59*). MION contrast agent was injected to increase signal-to-noise ratio. During fMRI, monkeys passively viewed blocks of faces and blocks of other objects to identify face-selective patches in the brain.

Before targeting fMRI-identified face patches with Neuropixels probes, we performed scout recordings with tungsten electrodes with 1 MΩ impedance (FHC) using grids designed with the software Planner (*60*). While inserting tungsten electrodes, we performed structural MRI scans to confirm correct targeting (Fig. 4a). Subsequently, we performed a total of 72 Neuropixels insertions. In order to perform very deep recordings (e.g., 42 mm from the craniotomy, Fig. 4d), we lowered a cannula holder to touch or gently push the dura. The cannula holder contained a short cannula to penetrate the dura. A probe holder, that held the probe, was slid through the cannula holder through matching dovetails. This dovetail mechanism was designed to ensure that the direction of probe movement matched the direction of the cannula, as even small differences in angles would risk breakage of the probe when inserted deeply into the cannula. The probe holder was advanced using an oil hydraulic micromanipulator (Narishige), but importantly, the precise direction of probe movement was constrained by the dovetail between probe holder and cannula holder rather than the micromanipulator.

Neuropixels data were recorded using SpikeGLX and OpenEphys, and spikes were sorted using Kilosort 3.0. To compute responses for Fig. 4c,e average spike rates from 50 ms to 250 ms after trial onset were computed, and baselines, averaged from 0 ms to 50 ms after trial onset, were subtracted.

### Functional connectivity via cross-correlation analysis

Functional interactions between pairs of neurons were measured with an established cross-correlation method. Cross-correlogram (CCG) were calculated using spike trains from pairs of simultaneously recorded neurons, either during the whole stimulus presentation period or during the inter-trial-intervals. The CCG is defined as:

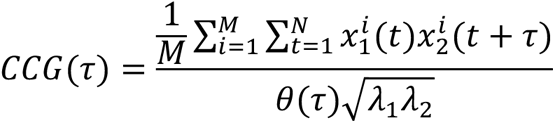

Where M is the number of trials, N is the number of time bins within a trial,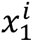 and 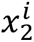 are the spike trains of neuron 1 and 2 from trial i, τ is the time lag relative to the reference spikes, and λ_1_and λ_2_ are the mean firing rate of the two neurons, respectively. θ(τ) is a triangular function calculated as θ(τ) = *N* − |τ| that corrects for the overlapping time bins at different time lags. A jitter corrected method was used to remove correlations caused by stimulus-locking or slow fluctuations.

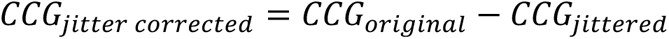

Where *CCG*_original_ and *CCG*_jittered_ are CCGs calculated with the above equation using original dataset and dataset with spike timing randomly perturbed (jittered) within the jitter window, respectively. The correction term (*CCG*_jittered_) captured slow correlation longer than the jitter window (caused by common stimulation or slow fluctuation in the population response), thus once it’s subtracted, only the fine temporal correlation is preserved. A 25-ms jitter window was chosen based on previous studies (*6*).

As with previous studies (*6, 11*), here a CCG is classified as significant if the peak of jitter-corrected CCG occurred within 10ms of zero time lag and if this peak is more than 7 standard deviations above the mean of the noise distribution (CCG flank).

### Current source density analysis and session alignment

For each session, we first flagged bad LFP channels by following the approach described in the International Brain Lab (IBL) spike sorting pipeline(*61*). Briefly, we identify dead channels, with unusually low similarity to the common average reference (CAR) signal, and noisy channels with high spectral power above 0.8 of the Nyquist frequency or low similarity with the high pass filtered (CAR). We low-pass filtered the LFP using a zero-phase, fifth order Butterworth filter with 25 Hz corner frequency. We also corrected in the frequency domain for the relative sampling offsets among the channels sharing a common ADC on the probe, as also described in the IBL pipeline. We then extracted LFP signals aligned to the onset of force production for each trial. We infilled the bad channels using kriging interpolation, using weights decreasing with distance *d* as ***w(d) ∝ e^−(d/d_0_)p^*** where *d_0_* = 20 μm and *p* = 1.3. We computed the event related potential (ERP) as the average of the aligned LFP signals across trials within each force profile condition and the average over all trials as well. We then resampled the ERP spatially along a vertical column running parallel with the center of the probe, sampled every 10 μm, again using kriging. To compute the current source density (CSD) for each condition individually and across all trials, we computed the second spatial derivative of the ERP using a smoothing, differentiating Savitzy Golay filter of second order, with a 450 μm frame size. To align the CSDs across sessions, we mean squared error as the loss function, i.e. normalized by the number of overlapping rows in the CSD. For each pair of the sessions, the loss function used the set of shared conditions collected in both experimental sessions and concatenated the CSDs for these shared conditions in time. If no conditions were present in both sessions, the grand average CSDs were used. We then found the optimal alignment jointly across all sessions by performing constrained optimization using a genetic algorithm.

---

**Fig. S1.**
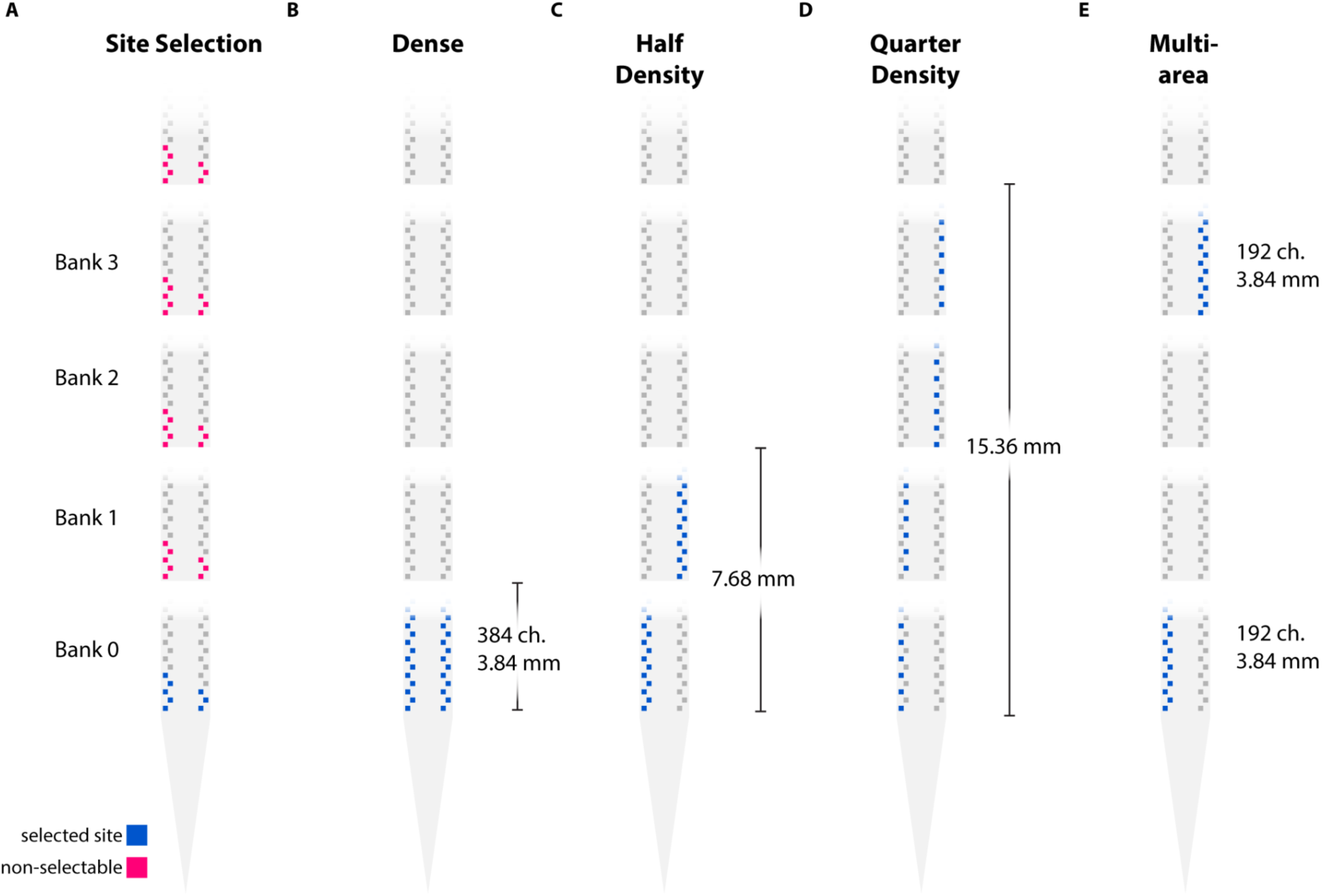
Site selection rules and common configurations. **(A)** Site selection rule - any electrode selected on any bank is unavailable for selection in other banks. Note that multiple electrodes can be connected to a single readout channel, as explored by (*2*), but this impacts the signal to noise and detection for small units, and is a specialized method and not currently in widespread use. **(B)** Dense recording from Bank 0, places 384 channels spanning 3.84 mm. **(C)** half density configuration covering Banks 0 and 1, spanning 7.68mm. **(D)** Quarter density, covering banks 0–3, spanning 15.36 mm. **(E)** multi-area recording.

**Fig. S2.**
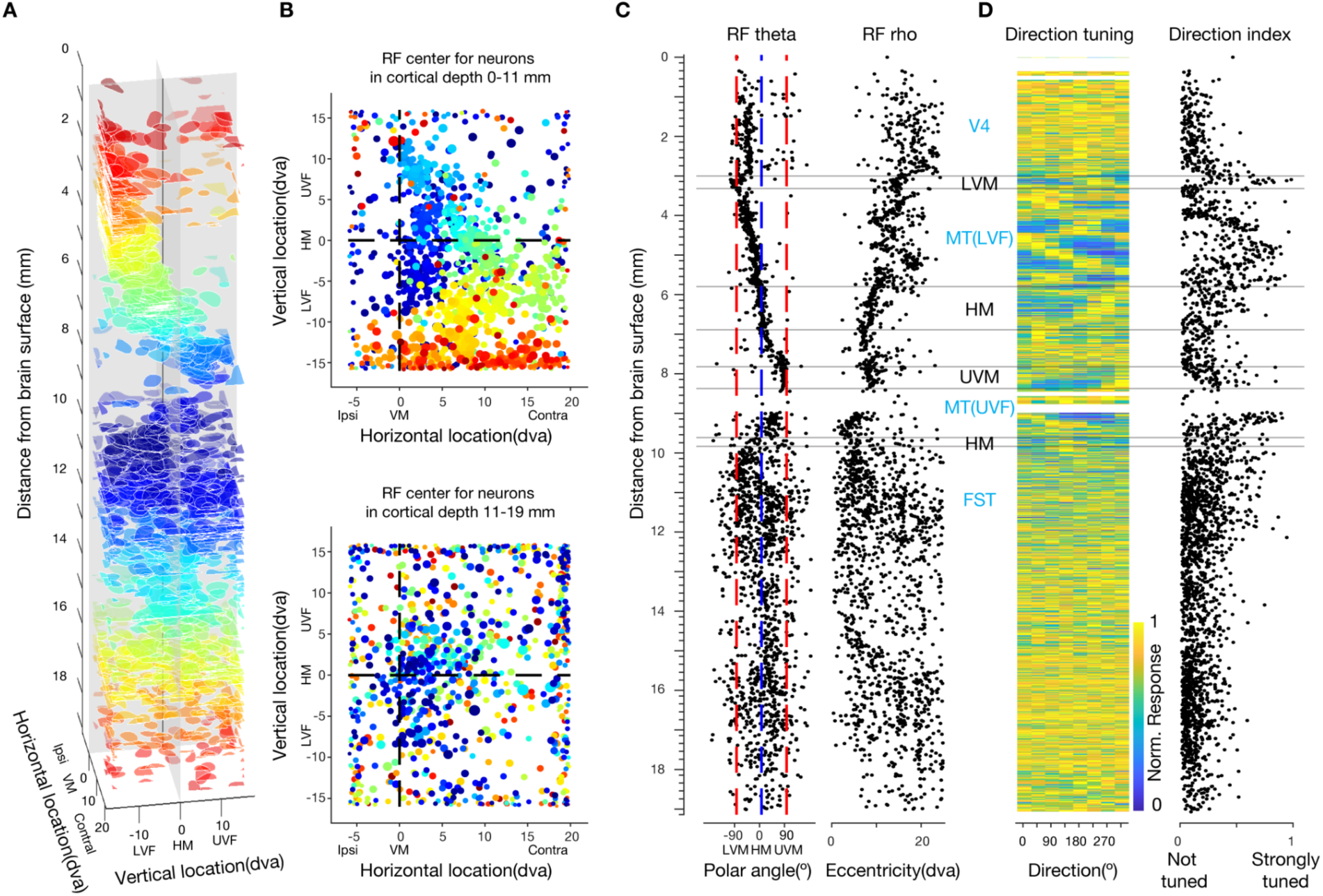
Retinotopic organization and functional properties of single neurons across multiple visual areas. **(A)** Distribution of receptive fields (RFs) of 2729 visually responsive neurons across cortical depth from the 5-banks recording across multiple visual areas. Color scale represents cortical depth. **(B)** Top-view of **A,** illustrating the progression of RFs across visual fields. RFs from the superficial and deeper part of the brain are demonstrated separately for clarity. **(C)** Polar angle (theta) and Eccentricity (rho) of each RF’s geometric center across cortical depth. **(D)** Left, heat map of evoked responses across drift directions of grating (vertical thickness is greater for less dense neuronal population). Color scale represents the magnitude of evoked responses. Right, direction index as quantified by the differences of responses to the preferred and its opposite direction divided by the sum of the two. In **C** and **D**, each neuron is plotted at its corresponding cortical depth. Horizontal lines denote the section of cortex where the center of RFs falls on Lower vertical meridians (LVM), horizontal meridian (HM), upper vertical meridian (UVM), and horizontal meridian (HM), respectively, superficial to deep. Putative visual areas are identified and labeled. LVF: lower visual field; UVF: upper visual field; FST: fundus of the superior temporal (FST) area.

**Fig. S3.**
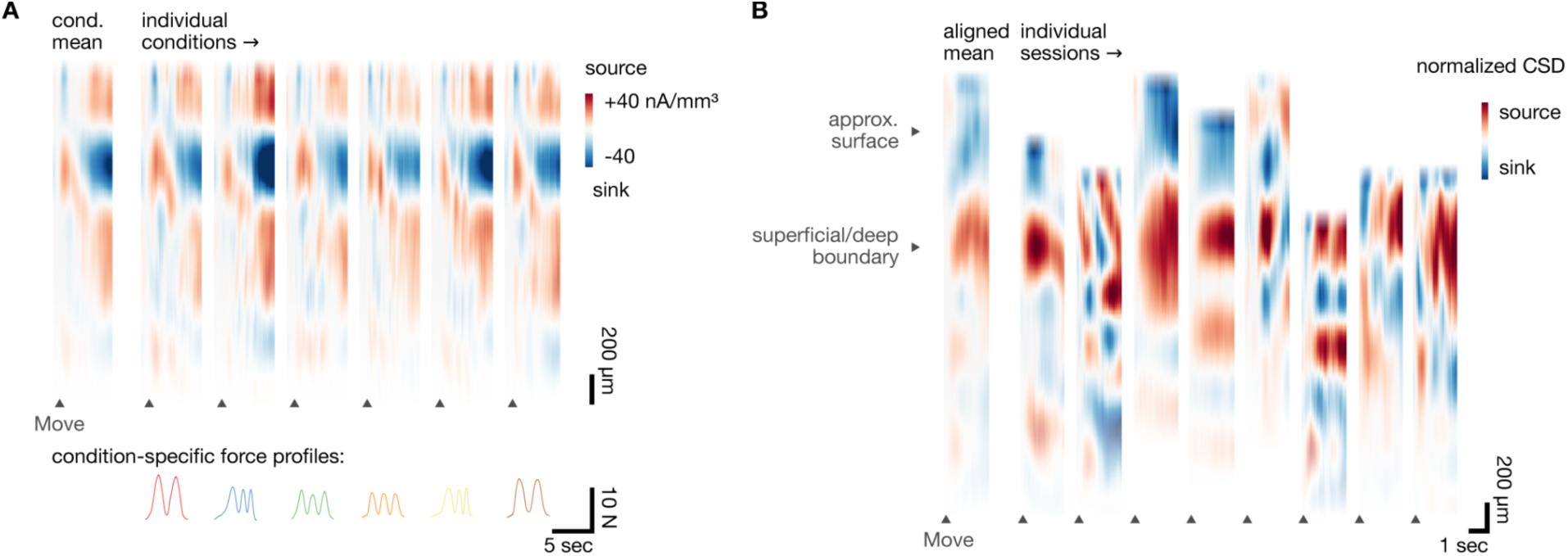
Aligning recordings to cortical lamina using current source density. **(A)** Single session current source density (CSD) plot during motor behavior. (top left) Mean CSD plot over all behavioral conditions, time locked to movement onset. (top right panels) Single-condition CSD plots illustrating spatial and temporal dynamics of CSD during behavior. (bottom row) Trial-averaged force behavior during pacman task for each condition. **(B)** Probe locations from Individual recording sessions aligned using CSD to infer recording depth relative to other sessions.

**Fig. S4.**
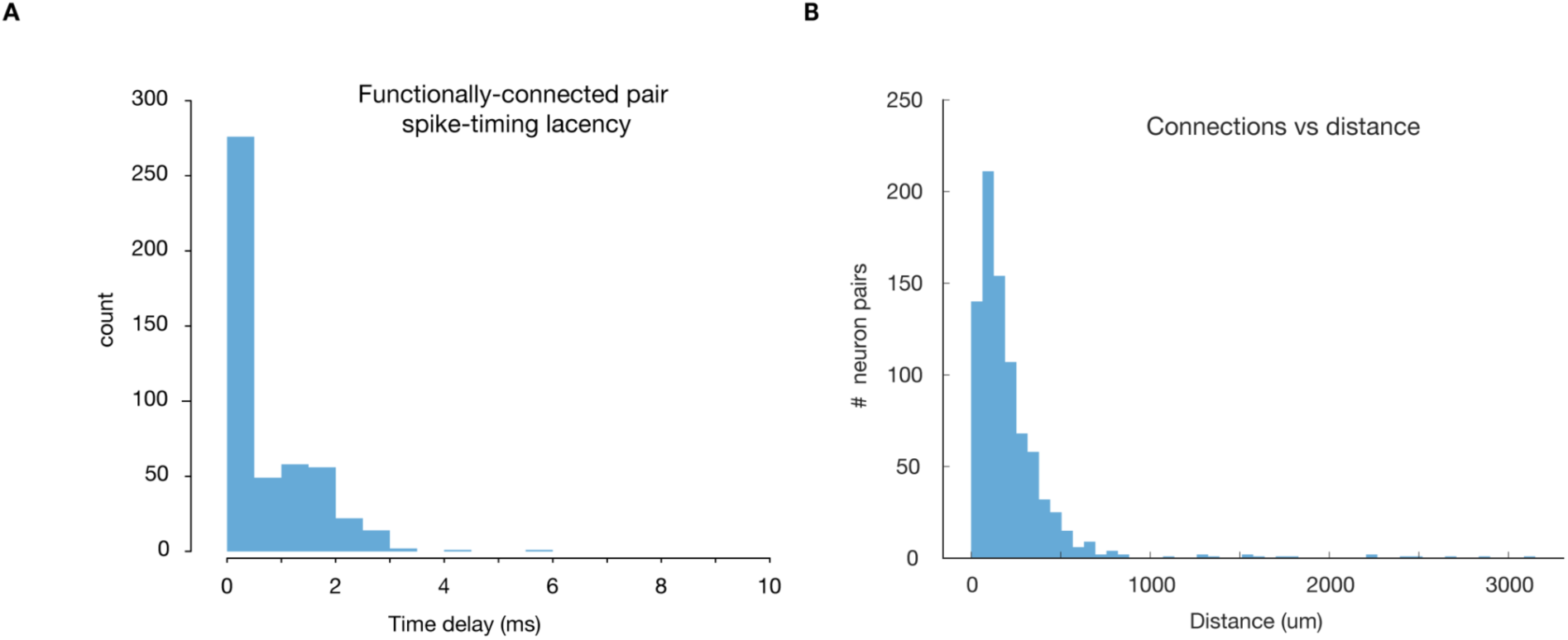
Functional connectivity metrics. **(A)** Latency distribution between identified peaks in cross-correlograms of neurons with statistically significant spike timing relationships. Example session, monkey C. **(B)** Number of functionally connected pairs of neurons as a function of the distance between two neurons.

**Fig. S5.**
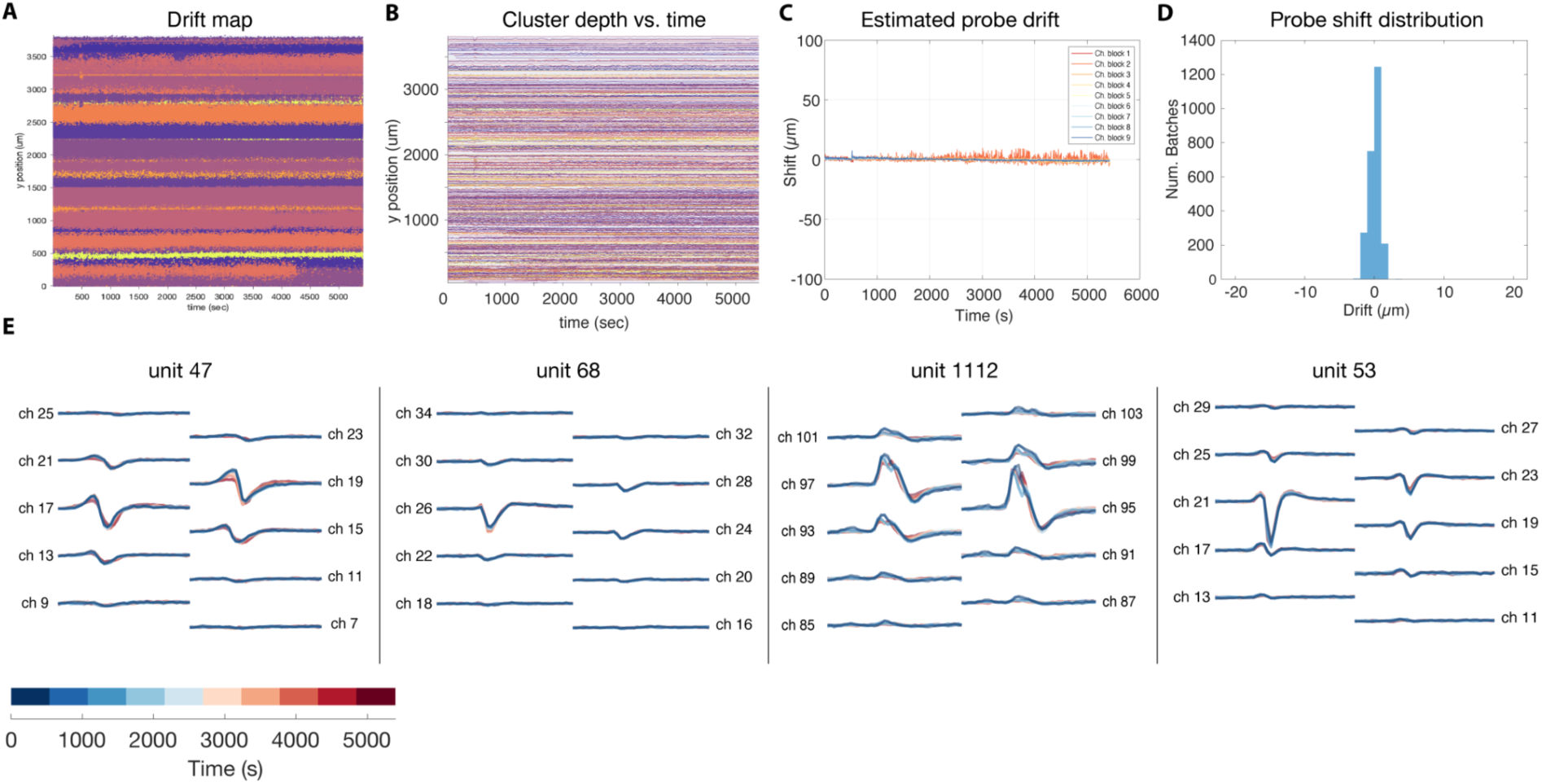
Acute recording stability. **(A)** Drift map of individual spikes, colored according to clusters identified via Kilosort 2.5. **(B)** Depth estimate of each cluster over time within a recording session. **(C)** Tissue drift relative to the probe estimated using Kilosort 2.5. Different traces indicate different blocks of channels on the probe. **(D)** Distribution of probe drift estimates across all alignment batches. **(E)** Neuron waveforms for four units identified using Kilosort 2.5. Individual traces represent median waveform calculated using 100 randomly spikes during one of 10 blocks during the session, with block indicated by line color. Data for panels **A-E** collected from Monkey I in the motor cortex during a motor behavioral experiment.

## References

1. J. J. Jun, N. A. Steinmetz, J. H. Siegle, D. J. Denman, M. Bauza, B. Barbarits, A. K. Lee, C. A. Anastassiou, A. Andrei, Ç. Aydın, M. Barbic, T. J. Blanche, V. Bonin, J. Couto, B. Dutta, S. L. Gratiy, D. A. Gutnisky, M. Häusser, B. Karsh, P. Ledochowitsch, C. M. Lopez, C. Mitelut, S. Musa, M. Okun, M. Pachitariu, J. Putzeys, P. D. Rich, C. Rossant, W.-L. Sun, K. Svoboda, M. Carandini, K. D. Harris, C. Koch, J. O’Keefe, T. D. Harris, Fully integrated silicon probes for high-density recording of neural activity. Nature. 551, 232–236 (2017).

2. N. A. Steinmetz, C. Aydin, A. Lebedeva, M. Okun, M. Pachitariu, M. Bauza, M. Beau, J. Bhagat, C. Böhm, M. Broux, S. Chen, J. Colonell, R. J. Gardner, B. Karsh, F. Kloosterman, D. Kostadinov, C. Mora-Lopez, J. O’Callaghan, J. Park, J. Putzeys, B. Sauerbrei, R. J. J. van Daal, A. Z. Vollan, S. Wang, M. Welkenhuysen, Z. Ye, J. T. Dudman, B. Dutta, A. W. Hantman, K. D. Harris, A. K. Lee, E. I. Moser, J. O’Keefe, A. Renart, K. Svoboda, M. Häusser, S. Haesler, M. Carandini, T. D. Harris, Neuropixels 2.0: A miniaturized high-density probe for stable, long-term brain recordings. Science. 372 (2021), doi:10.1126/science.abf4588.

3. N. A. Steinmetz, P. Zatka-Haas, M. Carandini, K. D. Harris, Distributed coding of choice, action and engagement across the mouse brain. Nature. 576, 266–273 (2019).

4. A. J. Peters, J. M. J. Fabre, N. A. Steinmetz, K. D. Harris, M. Carandini, Striatal activity topographically reflects cortical activity. Nature. 591, 420–425 (2021).

5. X. Jia, J. H. Siegle, C. Bennett, S. D. Gale, D. J. Denman, C. Koch, S. R. Olsen, High-density extracellular probes reveal dendritic backpropagation and facilitate neuron classification. J. Neurophysiol. 121, 1831–1847 (2019).

6. J. H. Siegle, X. Jia, S. Durand, S. Gale, C. Bennett, N. Graddis, G. Heller, T. K. Ramirez, H. Choi, J. A. Luviano, P. A. Groblewski, R. Ahmed, A. Arkhipov, A. Bernard, Y. N. Billeh, D. Brown, M. A. Buice, N. Cain, S. Caldejon, L. Casal, A. Cho, M. Chvilicek, T. C. Cox, K. Dai, D. J. Denman, S. E. J. de Vries, R. Dietzman, L. Esposito, C. Farrell, D. Feng, J. Galbraith, M. Garrett, E. C. Gelfand, N. Hancock, J. A. Harris, R. Howard, B. Hu, R. Hytnen, R. Iyer, E. Jessett, K. Johnson, I. Kato, J. Kiggins, S. Lambert, J. Lecoq, P. Ledochowitsch, J. H. Lee, A. Leon, Y. Li, E. Liang, F. Long, K. Mace, J. Melchior, D. Millman, T. Mollenkopf, C. Nayan, L. Ng, K. Ngo, T. Nguyen, P. R. Nicovich, K. North, G. K. Ocker, D. Ollerenshaw, M. Oliver, M. Pachitariu, J. Perkins, M. Reding, D. Reid, M. Robertson, K. Ronellenfitch, S. Seid, C. Slaughterbeck, M. Stoecklin, D. Sullivan, B. Sutton, J. Swapp, C. Thompson, K. Turner, W. Wakeman, J. D. Whitesell, D. Williams, A. Williford, R. Young, H. Zeng, S. Naylor, J. W. Phillips, R. C. Reid, S. Mihalas, S. R. Olsen, C. Koch, Survey of spiking in the mouse visual system reveals functional hierarchy. Nature. 592, 86–92 (2021).

7. W. E. Allen, M. Z. Chen, N. Pichamoorthy, R. H. Tien, M. Pachitariu, L. Luo, K. Deisseroth, Thirst regulates motivated behavior through modulation of brainwide neural population dynamics. Science. 364, 253 (2019).

8. S. Vesuna, I. V. Kauvar, E. Richman, F. Gore, T. Oskotsky, C. Sava-Segal, L. Luo, R. C. Malenka, J. M. Henderson, P. Nuyujukian, J. Parvizi, K. Deisseroth, Deep posteromedial cortical rhythm in dissociation. Nature. 586, 87–94 (2020).

9. R. J. Gardner, E. Hermansen, M. Pachitariu, Y. Burak, N. A. Baas, B. A. Dunn, M.-B. Moser, E. I. Moser, Toroidal topology of population activity in grid cells. Nature. 602, 123–128 (2022).

10. E. M. Trautmann, S. D. Stavisky, S. Lahiri, K. C. Ames, M. T. Kaufman, D. J. O’Shea, S. Vyas, X. Sun, S. I. Ryu, S. Ganguli, K. V. Shenoy, Accurate Estimation of Neural Population Dynamics without Spike Sorting. Neuron. 103, 292–308.e4 (2019).

11. E. B. Trepka, S. Zhu, R. Xia, X. Chen, T. Moore, Functional interactions among neurons within single columns of macaque V1. Elife. 11 (2022), doi:10.7554/eLife.79322.

12. X. Sun, D. J. O’Shea, M. D. Golub, E. M. Trautmann, S. Vyas, S. I. Ryu, K. V. Shenoy, Cortical preparatory activity indexes learned motor memories. Nature. 602, 274–279 (2022).

13. D. J. O’Shea, L. Duncker, W. Goo, X. Sun, S. Vyas, E. M. Trautmann, I. Diester, C. Ramakrishnan, K. Deisseroth, M. Sahani, K. V. Shenoy, Direct neural perturbations reveal a dynamical mechanism for robust computation. bioRxiv (2022), p. 2022.12.16.520768.

14. A. C. Paulk, Y. Kfir, A. R. Khanna, M. L. Mustroph, E. M. Trautmann, D. J. Soper, S. D. Stavisky, M. Welkenhuysen, B. Dutta, K. V. Shenoy, L. R. Hochberg, R. M. Richardson, Z. M. Williams, S. S. Cash, Large-scale neural recordings with single neuron resolution using Neuropixels probes in human cortex. Nat. Neurosci. 25, 252–263 (2022).

15. J. E. Chung, K. K. Sellers, M. K. Leonard, L. Gwilliams, D. Xu, M. E. Dougherty, V. Kharazia, S. L. Metzger, M. Welkenhuysen, B. Dutta, E. F. Chang, High-density single-unit human cortical recordings using the Neuropixels probe. Neuron (2022), doi:10.1016/j.neuron.2022.05.007.

16. E. M. Maynard, C. T. Nordhausen, R. A. Normann, The Utah intracortical Electrode Array: a recording structure for potential brain-computer interfaces. Electroencephalogr. Clin. Neurophysiol. 102, 228–239 (1997).

17. S. Musallam, M. J. Bak, P. R. Troyk, R. A. Andersen, A floating metal microelectrode array for chronic implantation. J. Neurosci. Methods. 160, 122–127 (2007).

18. D. A. Schwarz, M. A. Lebedev, T. L. Hanson, D. F. Dimitrov, G. Lehew, J. Meloy, S. Rajangam, V. Subramanian, P. J. Ifft, Z. Li, A. Ramakrishnan, A. Tate, K. Z. Zhuang, M. A. L. Nicolelis, Chronic, wireless recordings of large-scale brain activity in freely moving rhesus monkeys. Nat. Methods. 11, 670–676 (2014).

19. N. M. Dotson, S. J. Hoffman, B. Goodell, C. M. Gray, A Large-Scale Semi-Chronic Microdrive Recording System for Non-Human Primates. Neuron. 96, 769–782.e2 (2017).

20. D. Mao, E. Avila, B. Caziot, J. Laurens, J. D. Dickman, D. E. Angelaki, Spatial modulation of hippocampal activity in freely moving macaques. Neuron. 109, 3521–3534.e6 (2021).

21. E. M. Trautmann, D. J. O’Shea, X. Sun, J. H. Marshel, A. Crow, B. Hsueh, S. Vesuna, L. Cofer, G. Bohner, W. Allen, I. Kauvar, S. Quirin, M. MacDougall, Y. Chen, M. P. Whitmire, C. Ramakrishnan, M. Sahani, E. Seidemann, S. I. Ryu, K. Deisseroth, K. V. Shenoy, Dendritic calcium signals in rhesus macaque motor cortex drive an optical brain-computer interface. Nat. Commun. 12, 3689 (2021).

22. A. Bollimunta, S. R. Santacruz, R. W. Eaton, P. S. Xu, J. H. Morrison, K. A. Moxon, J. M. Carmena, J. J. Nassi, Head-mounted microendoscopic calcium imaging in dorsal premotor cortex of behaving rhesus macaque. Cell Rep. 35, 109239 (2021).

23. J. P. Rominger, B. J. Lin, Seamless Stitching For Large Area Integrated Circuit Manufacturing. SPIE Proceedings (1988), doi:10.1117/12.968412.

24. B. J. Lin, “The Paths To Subhalf-Micrometer Optical Lithography” in Optical/Laser Microlithography (SPIE, 1988), vol. 0922, pp. 256–269.

25. M. Pachitariu, N. Steinmetz, S. Kadir, M. Carandini, H. K. D. Kilosort, realtime spike-sorting for extracellular electrophysiology with hundreds of channels. bioRxiv (2016).

26. D. J. Felleman, D. C. Van Essen, Distributed Hierarchical Processing in the Primate Cerebral Cortex. Cerebral Cortex. 1 (1991), pp. 1–47.

27. D. Y. Tsao, W. A. Freiwald, T. A. Knutsen, J. B. Mandeville, R. B. H. Tootell, Faces and objects in macaque cerebral cortex. Nat. Neurosci. 6, 989–995 (2003).

28. G. A. Orban, D. Van Essen, W. Vanduffel, Comparative mapping of higher visual areas in monkeys and humans. Trends Cogn. Sci. 8, 315–324 (2004).

29. R. Gattass, C. G. Gross, Visual topography of striate projection zone (MT) in posterior superior temporal sulcus of the macaque. J. Neurophysiol. 46, 621–638 (1981).

30. W. M. Maguire, J. S. Baizer, Visuotopic organization of the prelunate gyrus in rhesus monkey. J. Neurosci. 4, 1690–1704 (1984).

31. R. Gattass, A. P. Sousa, C. G. Gross, Visuotopic organization and extent of V3 and V4 of the macaque. Journal of neuroscience. 8, 1831–1845 (1988).

32. R. P. Dum, P. L. Strick, Motor areas in the frontal lobe of the primate. Physiol. Behav. 77, 677–682 (2002).

33. J.-A. Rathelot, P. L. Strick, Subdivisions of primary motor cortex based on cortico-motoneuronal cells. Proc. Natl. Acad. Sci. U. S. A. 106, 918–923 (2009).

34. P. L. Strick, R. P. Dum, J.-A. Rathelot, The Cortical Motor Areas and the Emergence of Motor Skills: A Neuroanatomical Perspective. Annu. Rev. Neurosci. (2021), doi:10.1146/annurev-neuro-070918-050216.

35. S. H. Scott, The computational and neural basis of voluntary motor control and planning. Trends Cogn. Sci. 16, 541–549 (2012).

36. N. J. Marshall, J. I. Glaser, E. M. Trautmann, E. A. Amematsro, S. M. Perkins, M. N. Shadlen, L. F. Abbott, J. P. Cunningham, M. M. Churchland, Flexible neural control of motor units. bioRxiv (2022), p. 2021.05.05.442653.

37. C. Nicholson, J. A. Freeman, Theory of current source-density analysis and determination of conductivity tensor for anuran cerebellum. J. Neurophysiol. 38, 356–368 (1975).

38. U. Mitzdorf, Current source-density method and application in cat cerebral cortex: investigation of evoked potentials and EEG phenomena. Physiol. Rev. 65, 37–100 (1985).

39. D. Kobak, W. Brendel, C. Constantinidis, C. E. Feierstein, A. Kepecs, Z. F. Mainen, X.-L. Qi, R. Romo, N. Uchida, C. K. Machens, Demixed principal component analysis of neural population data. Elife. 5 (2016), doi:10.7554/eLife.10989.

40. J. D. Semedo, A. Zandvakili, C. K. Machens, B. M. Yu, A. Kohn, Cortical Areas Interact through a Communication Subspace. Neuron. 102, 249–259.e4 (2019).

41. S. Vyas, M. D. Golub, D. Sussillo, K. V. Shenoy, Computation Through Neural Population Dynamics. Annu. Rev. Neurosci. 43, 249–275 (2020).

42. P. Bao, L. She, M. McGill, D. Y. Tsao, A map of object space in primate inferotemporal cortex. Nature. 583, 103–108 (2020).

43. D. Y. Tsao, S. Moeller, W. A. Freiwald, Comparing face patch systems in macaques and humans. Proc. Natl. Acad. Sci. U. S. A. 105, 19514–19519 (2008).

44. J. K. Hesse, D. Y. Tsao, The macaque face patch system: a turtle’s underbelly for the brain. Nat. Rev. Neurosci. 21, 695–716 (2020).

45. L. Chang, D. Y. Tsao, The Code for Facial Identity in the Primate Brain. Cell. 169, 1013–1028.e14 (2017).

46. J. K. Hesse, D. Y. Tsao, A new no-report paradigm reveals that face cells encode both consciously perceived and suppressed stimuli. Elife. 9 (2020), doi:10.7554/eLife.58360.

47. M. N. Shadlen, R. Kiani, Decision making as a window on cognition. Neuron. 80, 791–806 (2013).

48. R. Ratcliff, G. McKoon, The diffusion decision model: theory and data for two-choice decision tasks. Neural Comput. 20, 873–922 (2008).

49. N. A. Steinemann, G. M. Stine, E. M. Trautmann, A. Zylberberg, D. M. Wolpert, M. N. Shadlen, Direct observation of the neural computations underlying a single decision. bioRxiv (2022), p. 2022.05.02.490321.

50. G. M. Stine, E. M. Trautmann, D. Jeurissen, M. N. Shadlen, A neural mechanism for terminating decisions. bioRxiv (2022), p. 2022.05.02.490327.

51. X. Jia, J. H. Siegle, S. Durand, G. Heller, T. K. Ramirez, C. Koch, S. R. Olsen, Multi-regional module-based signal transmission in mouse visual cortex. Neuron. 110, 1585–1598.e9 (2022).

52. A. Kohn, M. A. Smith, Stimulus dependence of neuronal correlation in primary visual cortex of the macaque. J. Neurosci. 25, 3661–3673 (2005).

53. G. C. DeAngelis, G. M. Ghose, I. Ohzawa, R. D. Freeman, Functional micro-organization of primary visual cortex: receptive field analysis of nearby neurons. J. Neurosci. 19, 4046–4064 (1999).

54. D. Y. Ts’o, C. D. Gilbert, T. N. Wiesel, Relationships between horizontal interactions and functional architecture in cat striate cortex as revealed by cross-correlation analysis. J. Neurosci. 6, 1160–1170 (1986).

55. C. Windolf, A. C. Paulk, Y. Kfir, E. Trautmann, S. Garcia, D. Meszéna, W. Muñoz, I. Caprara, M. Jamali, J. Boussard, Z. M. Williams, S. S. Cash, L. Paninski, E. Varol, Robust Online Multiband Drift Estimation in Electrophysiology Data. bioRxiv (2022), p. 2022.12.04.519043.

56. E. K. Lee, H. Balasubramanian, A. Tsolias, S. U. Anakwe, M. Medalla, K. V. Shenoy, C. Chandrasekaran, Non-linear dimensionality reduction on extracellular waveforms reveals cell type diversity in premotor cortex. eLife. 10 (2021), doi:10.7554/elife.67490.

57. B. Karsh, Others, SpikeGLX (2016).

58. N. J. Marshall, J. I. Glaser, E. M. Trautmann, E. A. Amematsro, S. M. Perkins, M. N. Shadlen, L. F. Abbott, J. P. Cunningham, M. M. Churchland, Flexible neural control of motor units. Nat. Neurosci. 25, 1492–1504 (2022).

59. D. Y. Tsao, W. A. Freiwald, R. B. H. Tootell, M. S. Livingstone, A cortical region consisting entirely of face-selective cells. Science. 311, 670–674 (2006).

60. S. Ohayon, D. Y. Tsao, MR-guided stereotactic navigation. J. Neurosci. Methods. 204, 389–397 (2012).

61. International Brain Laboratory, K. Banga, J. Boussard, G. A. Chapuis, M. Faulkner, K. D. Harris, J. M. Huntenburg, C. Hurwitz, H. D. Lee, L. Paninski, C. Rossant, N. Roth, N. A. Steinmetz, C. Windolf, O. Winter, “Spike sorting pipeline for the International Brain Laboratory” (2022), doi:10.6084/m9.figshare.19705522.v1.

